# Sequestration of SerRS through LLPS Impairs Localized Translation and Contributes to Antibiotic Persistence

**DOI:** 10.1101/2024.12.31.630848

**Authors:** Ziyin Zhang, Daqian Li, Bo Zheng, Jiafeng Liu

**Author notes:** Correspondence to Jiafeng Liu.

## Abstract

Antibiotic-tolerant persisters contribute to the emergence of resistance, posing a significant challenge to the efficacy of antibiotic therapies. Despite extensive research, the mechanisms underlying persistence remain inadequately understood. By tracking the evolution of exponential-phase bacterial populations subjected to intermittent high-dose ertapenem exposure, we characterized the evolved strains in terms of tolerance. Mutant strains, harboring mutations in the seryl-tRNA synthetase gene (*serS)*, exhibited abrupt growth arrest upon serine depletion during exponential growth, resembling the persistence phenotype induced by serine hydroxamate (SHX). Under serine starvation, the mutated SerRS protein was sequestrated into liquid-liquid phase separation (LLPS)-driven condensates, disrupting their composition and impairing localized translation. This event precipitated growth arrest and dormancy in the *SerS*^*T*^ strain, triggering persistence. Our findings reveal an unrecognized role for aminoacyl-tRNA synthetases (aaRSs) in modulating bacterial condensates and provide insights into the molecular mechanisms underlying bacterial persistence.

**Graphic abstract:** SerRS Recruitment Disrupts Localized Translation in DeaD-marked Condensates. Upon serine starvation, SerRS^T^ partitions into LLPS-driven DeaD-marked condensates, impairing their localized translation activity and suggesting mechanisms underlying persistence arising during exponential phase. These evolutionarily conserved condensates orchestrate a robust translation program to enable stress responses and instruct cell fate decisions in bacterial populations.

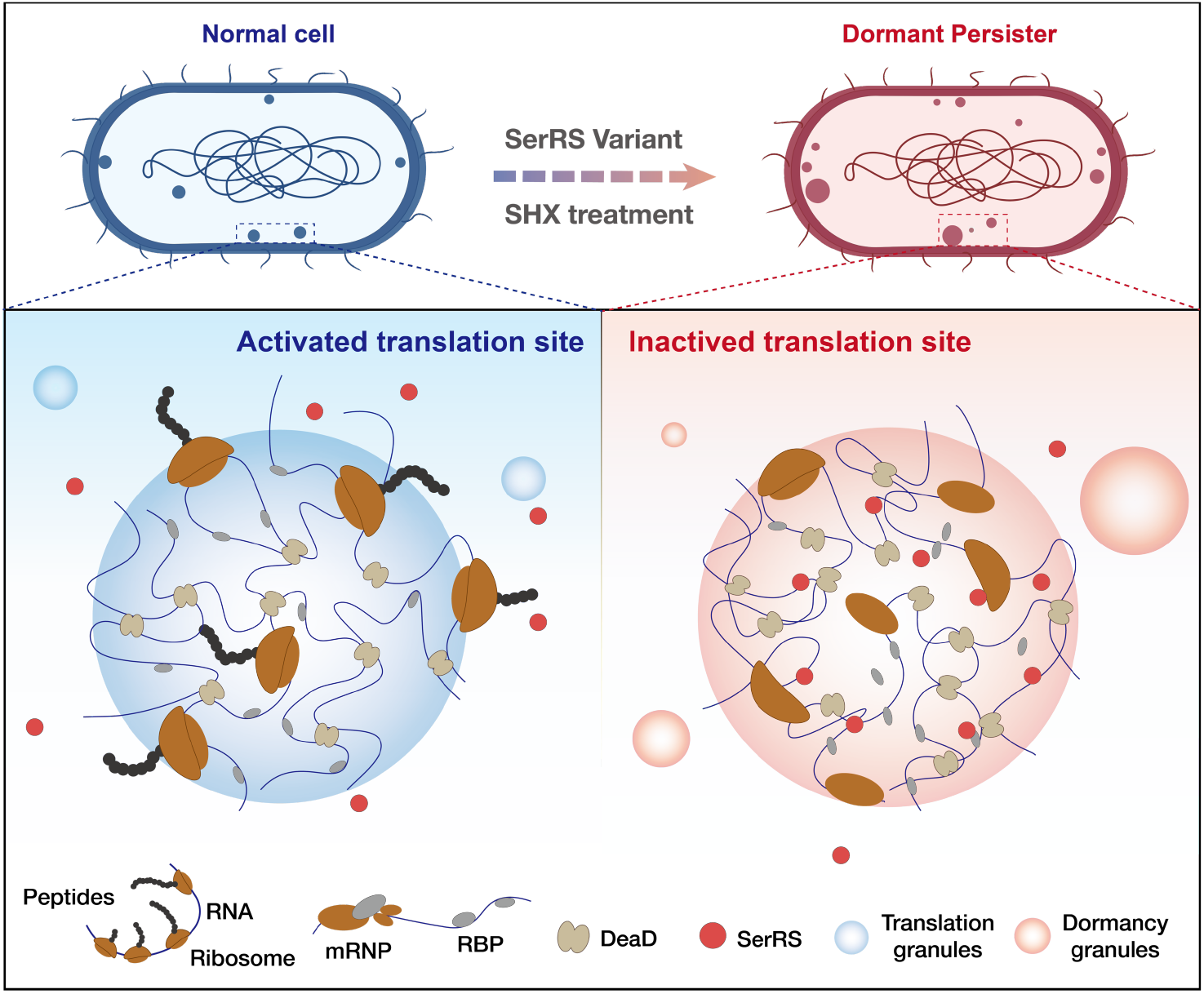

## Introduction

Bacteria persistence refers to a subpopulation of tolerant bacteria that can survive intensive antibiotic treatment without being resistant, a phenomenon termed heterotolerance (*1*). In clinical settings, persistence contributes to chronic infections and antibiotic treatment failure (*2*– *5*). By surviving high-dose antibiotics, persisters also serve as reservoirs for resistance evolution (*6*–*8*). Triggered persistence, induced by nutrient starvation during the stationary phase, delays regrowth upon transfer to fresh medium. By extending the lag phase of dormancy, non-dividing cells exhibit enhanced survival under antibiotic treatment (*1, 9, 10*). While mutations in aaRS and related genes have been reported to modulate triggered persistence, the underlying mechanism remain elusive (*7, 11*–*16*). Persistence can also arise spontaneously during exponential growth, wherein a small subpopulation enters a dormant state as an altruistic strategy within the actively growing population (*4, 10, 17*), yet the mechanism involved remain poorly understood. We therefore tested whether a distinct type of persister could evolve from an exponential-phase population under cyclic antibiotic treatment, and further investigated the mechanisms underlying persistence formation.

## Results

### Lag-time independent tolerance

In mid-exponential phase, *E. coli* exhibited a pronounced increase in persister-cell formation (Fig. S1a) (*17*). To investigate this persistence, we followed the evolution of log-phase populations of *E. coli* KLY under periodical exposure to a high concentration of ertapenem (2 μg/mL, approximately 100 times the minimum inhibitory concentration (MIC), Fig. S1b, Fig. 1a). Over several cycles, the killing efficiency of ertapenem declined by over two orders of magnitude (Fig. 1b). Critically, the MICs of ertapenem for clones isolated from the evolved lines remained indistinguishable from that for the ancestor, indicating that the increased survival was due to tolerance rather than resistance (Fig. 1c).

**Fig. 1.**
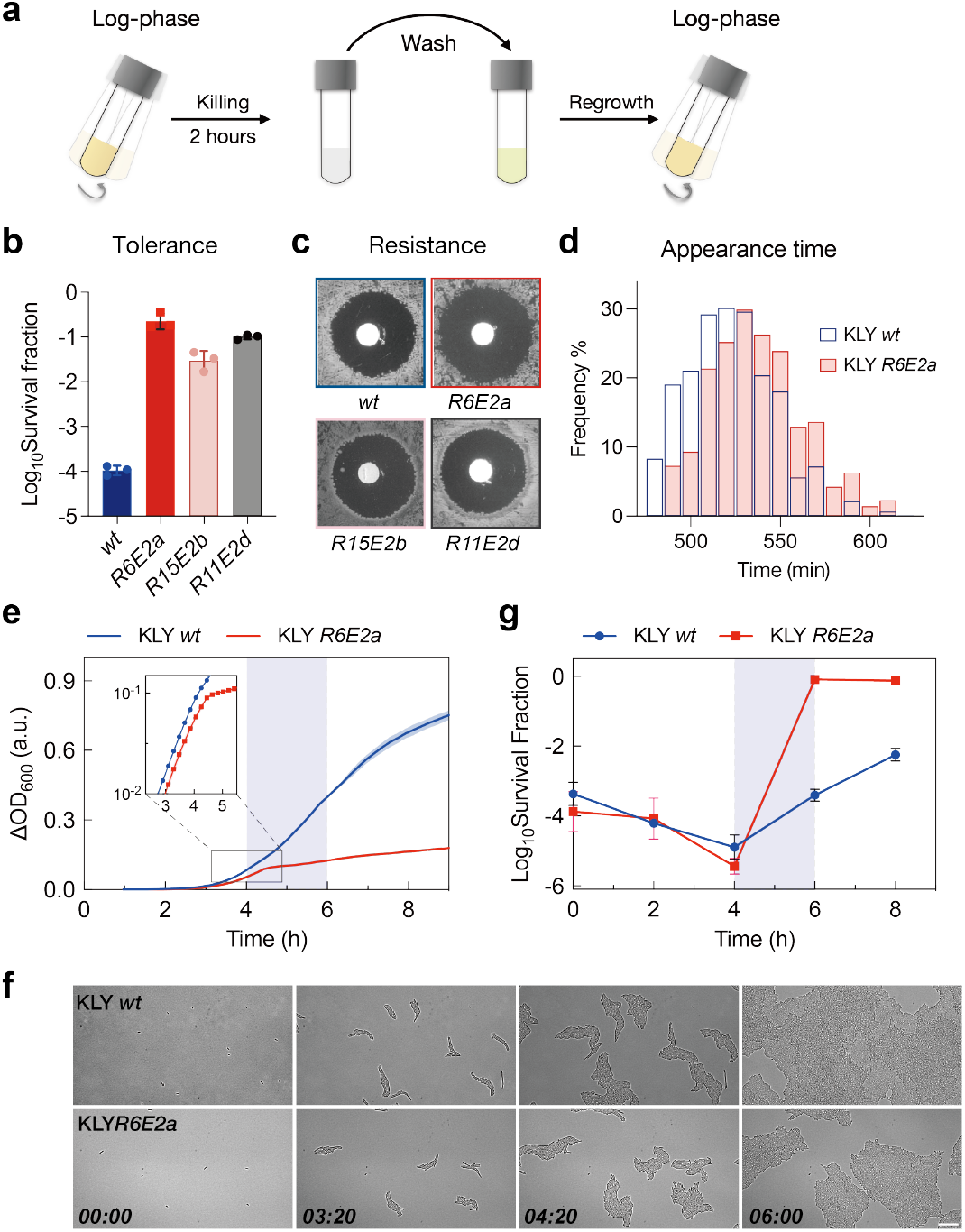
A lag-independent persistence mediated by growth arrest. (a) Experimental design for cyclic antibiotic exposure. In each cycle, log-phase cultures were treated with 2 μg/mL ertapenem for 2 hours. The cultures were then washed to remove the drug, resuspended in fresh LB media, and allowed to regrow to the exponential phase (OD_600_ ≈ 0.12). (b) Survival of the evolved and ancestral strains after 2 μg/mL ertapenem treatment for 3 hours. Data are presented as the mean ± s.d. from three independent experiments. The clones *R6E2a, R15E2b* and *R11E2d* were isolated from the 6^th^, 15^th^, and 11^th^ rounds of evolution, respectively. (c) MIC test performed using disc diffusion antibiotic sensitivity testing with 1 μg ertapenem. (d) Measurement of the lag-time distribution using the automated scanner system ScanLag. The histograms show the proportion of colony-forming units (CFUs) detected at each time point. Mean lag times are 524.5 min (*wt*) and 540.1 min (*R6E2*a). Sample sizes are n = 189 (*wt*) and 180 (*R6E2*a). (e) Growth curves of the evolved *R6E2a* and the ancestral strains. Cultures were grown in a 96-well plate, and OD_600_ was measured over time. Inset: OD_600_ data plotted on a log scale showed an abrupt switch of growth rate in mutant strain. (f) Phase-contrast images from time-lapse microscopy of single cells from an exponential culture plated onto fresh LB medium, illustrating the sudden growth arrest of the evolved strain. Scale bar, 20 μm. Times are indicated in hours and minutes. (g) Dynamics of the persister fraction over the cell growth duration. Cultures were sampled at an interval of 2 hours for an antibiotic killing assay. The evolved *SerS*^*T*^ strain exhibited a rapid accumulation of persister cells between 4 and 6 hours of growth, compared to the slower rise observed in the wild-type strain.

A tolerant strain KLY*R6E2a*, first established in the sixth cycle of antibiotic treatment, was selected to be characterized. This strain exhibited a similar appearance-time distribution (Fig. 1d) and growth rate (Fig. 1e, inset) to the ancestor, indicating lag-time-independent tolerance. In contrast, the mutant displayed abrupt growth arrest during the mid-exponential phase (ΔOD_600_ ≈ 0.1, Fig. 1e). Microscopic observations revealed that both ancestor and mutant strains started division within half an hour; however, the mutant experienced a sudden replication slowdown, whereas the ancestor maintained continuous growth (Fig. 1f, Fig. S1c and Supplemental Movie-1). The increased persistence in KLY*R6E2a* correlated with the growth arrest (Fig. 1e,1g and S1a), consistent with the fact that ertapenem primarily targets rapidly dividing cells. Surprisingly, whole-genome sequencing of clones isolated from independent evolved populations identified distinct mutations in the *serS* gene (Table S1), encoding seryl-tRNA synthetase. Previous studies have implicated mutations in aaRS genes in mediating tolerance through extended lag time (*7, 11*–*13*). The identification of analogous aaRS mutations in a distinct tolerance phenotype (KLY*R6E2a, SerS*^*T*^) suggests a shared mechanism underlying persistence across diverse conditions.

### Serine starvation as a trigger of persistence

The growth arrest observed in the mutant strain KLY*R6E2a* (*SerS*^*T*^) was specifically restored by the addition of serine, the cognate amino acid for seryl-tRNA synthetase, whereas supplementation with other amino acids had no effect (Fig. S2a). Final cell densities correlated directly with serine concentration (Fig. 2a). Moreover, serine supplementation re-sensitized the *SerS*^*T*^ strain to antibiotic treatment (Fig. 2b), confirming that the growth-arrest induced tolerance was driven by serine deficiency.

**Fig. 2.**
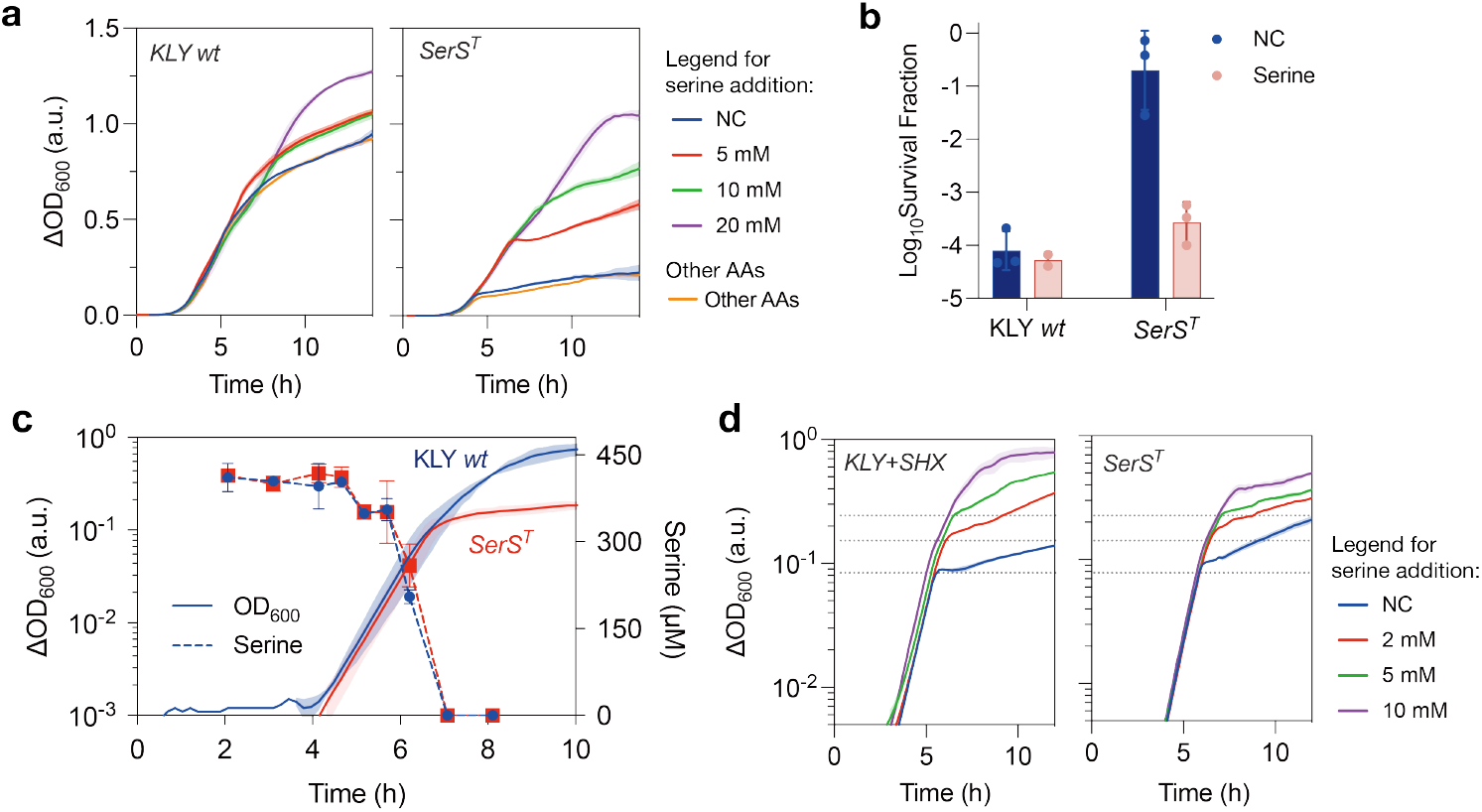
Serine-starvation triggered persistence. (a) Growth curves of the ancestral and *SerS*^*T*^ strains in LB medium with varying concentrations of serine added. (b) Survival of the ancestral and evolved strains after 2 μg/mL ertapenem treatment for 3 hours, with or without (NC) 2 mM serine supplementation in LB medium. Data are presented as the mean ± s.d. of three independent experiments. (c) Growth curves of the ancestral and tolerant *SerS*^*T*^ strains, along with the serine consumption profiles throughout the entire growth period. (d) Effect of SHX (1 mM) addition on cell growth of the ancestor KLY strain and its reversal by serine. Data are presented as the mean ± s.d. of three independent replicates.

Serine is preferentially utilized over other amino acids during bacterial growth, as previously reported (*18*–*20*). To investigate this further, we employed liquid chromatography-mass spectrometry (LC-MS) to monitor extracellular serine concentrations, revealing a sharp depletion during the exponential phase (Fig. 2c). Upon serine exhaustion, the tolerant strain shortly entered a dormant state, whereas the ancestral strain maintained growth (Fig. 2c). Despite the presence of alternative nutrients in the medium, *SerS*^*T*^ cells exhibited signs of serine starvation (Fig. S2b). Consistent results were obtained in the M9 medium supplemented with 20 amino acids (denoted 20AAs), where the *SerS*^*T*^ strain displayed a growth defect similar to that observed in LB medium (Fig. S2c), with sudden growth stasis alleviated by serine supplementation in a dose-dependent manner (Fig. S2d). Collectively, these results establish serine depletion as a critical factor driving the growth arrest in the *SerS*^*T*^ strain.

Serine hydroxamate (SHX), a serine analogue and competitive inhibitor of SerRS, induces serine starvation, triggering growth arrest and persistence in both wild-type and stringent response-relaxed mutant (*21*–*23*). SHX treatment caused immediate growth inhibition upon serine exhaustion in KLY, identical to the response observed in the *SerS*^*T*^ mutant (Fig. 2d, S2e). Additionally, both SHX-treated KLY and *SerS*^*T*^ strains were equally rescued by serine supplementation at varying concentrations (Fig. 2d). These findings suggest that serine starvation could be a common driver of persistence, whether it is triggered by SHX treatment or mutations in SerRS. Since the wild-type KLY (*relA1, spoT1*) is a relaxed strain (*24, 25*), this serine starvation-induced persistence in the *SerS*^*T*^ strain appears to be independent of (p)ppGpp mediated stringent response (*21, 26, 27*).

### The *SerS*^*T*^ strain exhibits prolonged dormancy in amino-acid free M9 medium

Upon transfer to amino acid-free M9 medium, the *SerS*^*T*^ mutant exhibited significantly prolonged lag times relative to KLY, although their growth rates were comparable (Fig. 3a). Interestingly, SHX treatment induced a similar extension of lag time in KLY strain (Fig. 3a, S3a). Time-lapse microscopy revealed that while ancestral KLY cells resumed division within four hours, a substantial fraction (∼60%) of *SerS*^*T*^ cells remained dormant even after 16 hours of incubation (Fig. 3b, 3c, and Supplemental Movie-2). Serine supplementation reduced the lag time in a dose-dependent manner (Fig. 3d, S3b), suggesting that the extended lag time observed in M9 medium was also attributable to serine starvation. This serine starvation-induced dormancy resembled the persistence with extended lag time reported in response to SHX exposure (*21*–*23*) or nutrient depletion in the stationary phase (*10, 28*).

**Fig. 3.**
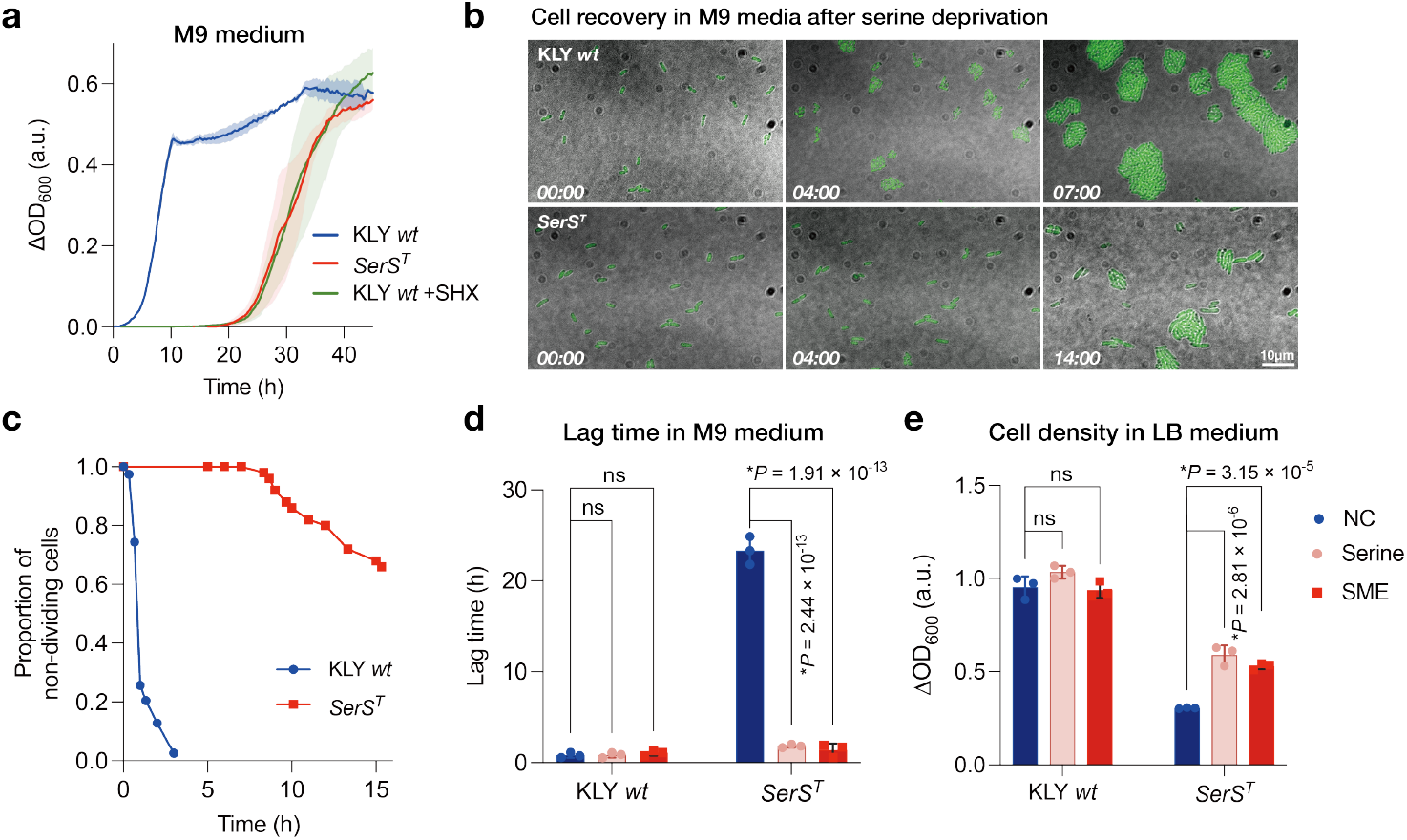
Supplementation of serine and analogues restored growth defect in the *SerS*^*T*^ strain. (a) Growth curves of the ancestor KLY and *SerS*^*T*^ strains, as well as the ancestor treated with SHX (1 mM), were generated from cell cultures initially grown in LB medium till ΔOD_600_ ≈ 0.1, followed by transfer into M9 medium via a 1:1000 dilution. (b) Time-lapse microscopy of single bacteria transferred from LB cultures to fresh M9 media, showing the extended lag time of the evolved strain. Time points are indicated in hours and minutes. Scale bar, 10 μm. (c) Proportion of non-dividing cells over incubation time in fresh M9 medium, related to (b). (d) Serine and its analogues significantly reduced the lag time in M9 minimal media. Serine, 0.1 mM; SME, Serine methyl ester, 0.1 mM. (e) Serine and its analogues restored the serine starvation-triggered growth arrest in LB medium. Serine, 5 mM; SME, 5 mM. Data are presented as the mean ± s.d., *P* values were calculated using two-tailed t-tests.

Although glycine and, to a limited extent, threonine can serve as metabolic substitutes for serine (*29*), neither alleviated the growth arrest observed in LB medium (Fig. S2a). SHX-induced persistence is canonically attributed to the competitive inhibition of SerRS, impairing tRNA^Ser^ aminoacylation and disrupting protein synthesis (*30, 31*). Notably, these serine analogues, including serine methyl ester (SME), serinamide and SHX, act as antimetabolites and competitive inhibitors of SerRS (*31*), paradoxically facilitating the exit of *SerS*^*T*^ cells from dormancy in M9 medium (Fig. S3a, S3c-S3d, 3d) or restoring growth arrest in LB medium as effectively as serine (Fig. 3e). These findings suggest that serine and its analogues might have roles that extend beyond their conventional functions as metabolites or tRNA substrates, potentially facilitating the exit from dormancy through interactions with the SerRS^T^ protein.

### SerRS^T^ preferentially translocates into DEAD-Box ATPase-marked condensates

Emerging evidence suggests that bacterial dormancy may involve protein condensates (*32*–*34*). In eukaryotic cells, ATP-dependent helicases, RNA-binding proteins, chaperonins, and aaRSs are conserved components of condensates implicated in various biological processes, such as degenerative disease, circadian clock regulation, and cell fate decisions (*35*–*37*). We hypothesize that the SerRS^T^ protein participates in biological condensates upon serine deprivation, thereby regulating cell growth dynamics (e.g., growth arrest in LB medium or extended lag time in M9 medium) and driving persistence.

To investigate the subcellular localization of SerRS, we used CRISPR-Cas9 to insert an mCherry tag into the endogenous *serS*^*wt*^ or *serS*^*T*^ loci in genomes, enabling real-time tracking in living cells. Microscopy observations revealed that SerRS^T^, but not the SerRS^wt^, preferentially localized to cell-pole granules during the early stationary phase (Fig. S4a). This finding was further corroborated by co-immunoprecipitation, which showed that SerRS^T^ was significantly enriched in the insoluble condensate fraction compared to SerRS^wt^ (Fig. 4a). We subsequently performed quantitative mass spectrometry to identify the insoluble proteome. Compared to the KLY wild-type strain, the abundance of SerRS^T^ was significantly increased in the insoluble pellet of the mutant strain (Fig. 4b). In addition to SerRS^T^, the RNA-dependent DEAD-box ATPase DeaD (encoded by the *deaD* gene) was also highly enriched in the insoluble pellets of the *SerS*^*T*^ strain (Fig. 4b, S4b). Since DEAD-box ATPases are conserved regulators of biological condensates across species (*38*–*41*), we hypothesize that DeaD protein might be involved in regulating the SerRS^T^-related condensates and growth defect.

**Fig. 4.**
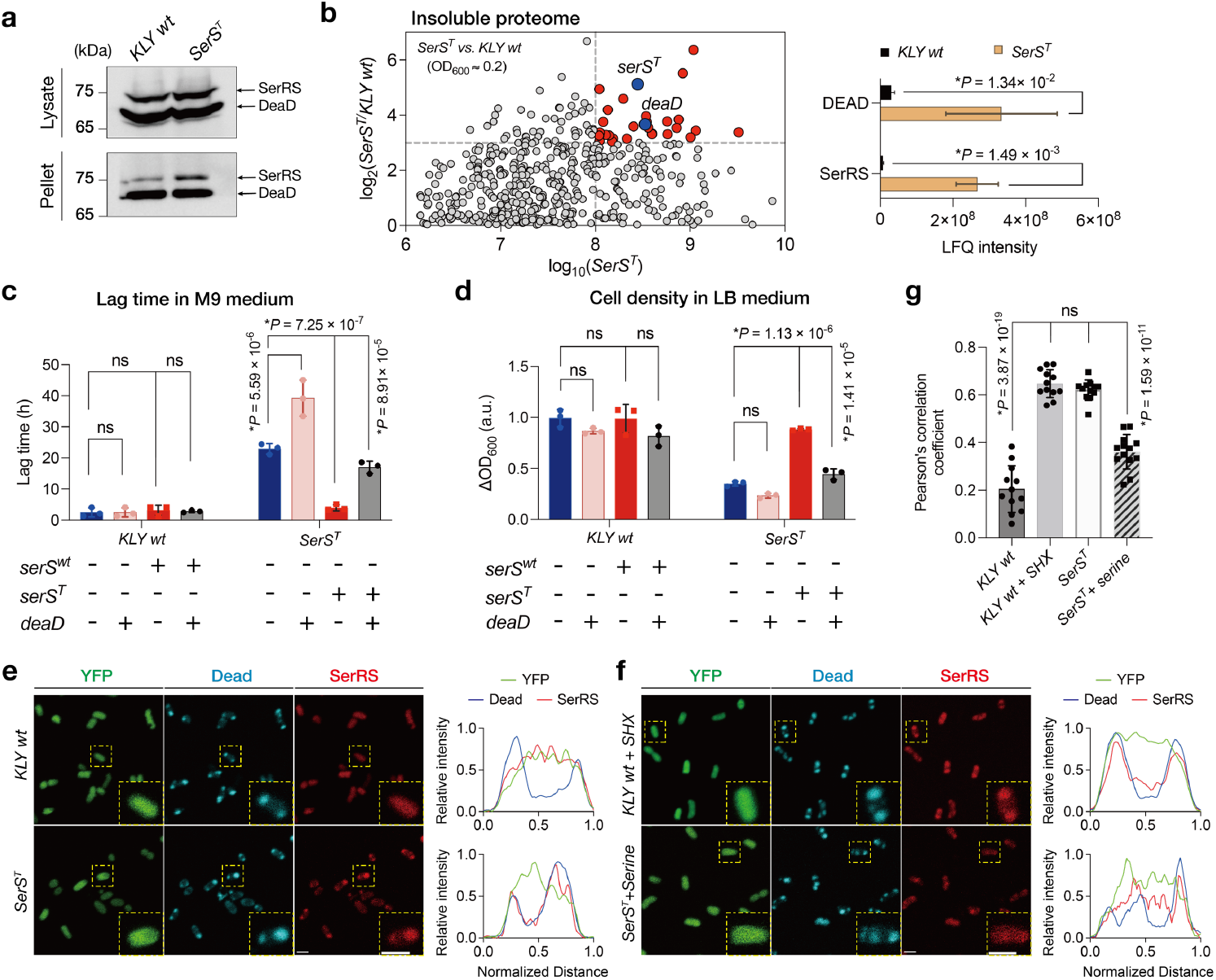
SerRS^T^ aberrantly translocated into the DeaD-condensates upon serine deprivation. (a) Immunoblotting for detecting endogenous SerRS in total cell lysate and pellet of the KLY *wt* and *SerS*^*T*^ strains. An mCherry tag was inserted into the endogenous *serS* loci in genomes, while His-tagged DeaD expressed from a plasmid and induced with tetracycline (0.025 μg/mL). (b) MS analysis of insoluble proteome in the *SerS*^*T*^ strain relative to the ancestor KLY control. Left panel, scatterplot showing the protein level changes. The y axis showed the log_2_ fold changes of the *SerS*^*T*^ strain over KLY strain, and x axis represented the log_2_ protein abundance of insoluble condensates in the mutant; right panel, label-free quantitation (LFQ) intensity of SerRS and DeaD in the insoluble pellets. Data are presented as the mean ± s.d. of three independent replicates, *P* values were calculated using two-tailed t-tests. (c-d) The lag time in M9 medium (c) and cell density in LB (d) were modulated by the overexpression of DeaD and SerRS. Inducer concentrations were 1 mM IPTG for DeaD expression under *P*_*Lac1*_ promoter, 0.05 μg/mL tetracycline for the expression of *serS*^*wt*^ or *serS*^*T*^ under the *P*_*tet*_ promoter. Data are presented as the mean ± s.d., *P* values were calculated using two-tailed t-tests. (e) Representative images of mCherry-tagged SerRS in cells expressing ECFP-tagged DeaD. (f) Representative images of mCherry-tagged SerRS in cells expressing ECFP-tagged DeaD with addition of 10 mM SHX in KLY *wt* cells and 10 mM serine in the *SerS*^*T*^ cells. Line scans show the related intensity profiles of SerRS^wt^ with DeaD in ancestral cells and SerRS^T^ with DeaD in mutated cells. Inset: higher magnification of the yellow boxed area. Scale bar, 2 μm. (g) Pearson’s correlation coefficient. Data are presented as the mean ± s.d., *P* values were calculated using two-tailed t-tests, sample sizes are n = 12 (KLY *wt*), 12 (KLY *wt* + SHX), 12 (*SerS*^*T*^) and 13 (*SerS*^*T*^ + Serine).

To test this hypothesis, we introduced a plasmid expressing DeaD into the *SerS*^*T*^ strain, resulting in a marked increase in lag time (Fig. 4c). Overexpression of the mutated *serS*^*T*^ gene alleviated the growth defect in LB medium and reduced the extended lag time in M9 medium under finely tuned inducer concentrations (Fig. 4c, 4d, S5a). However, the restored effect of *serS*^*T*^ overexpression was abolished by co-expression of DeaD protein, suggesting that both DeaD and SerRS^T^ function as regulators of growth dynamics with opposing effects (Fig. 4c, 4d, S5b). Knockout of *deaD* gene in the *SerS*^*T*^ tolerant strain minimally affected cell growth and lag time (Fig. S4c), potentially due to functional redundancy with other DEAD-box ATPases, such as SrmB and RhlE (*42*).

Fluorescence microscopy of the KLY and *SerS*^*T*^ strains, co-expressing mCherry-tagged SerRS (SerRS^wt^-mCherry or SerRS^T^-mCherry) and ECFP-tagged DeaD, revealed strong co-localization of SerRS^T^ with DeaD-marked condensates upon growth arrest, whereas SerRS^wt^ remained diffusely distributed in the cytoplasm (Fig. 4e, 4g). In contrast, both SerRS^wt^ and SerRS^T^ were evenly distributed throughout the cytoplasm prior to the growth stasis (Fig. S4d, S4e). Moreover, DEAD-box helicases SrmB and RhlE co-localized with DeaD, supporting that these helicases, together with DeaD, could be redundant modelers of condensates involved in modulating dormancy (Fig. S4f-4i).

Given that SHX treatment induced a growth defect comparable to the *SerS*^*T*^ strain, we further examined whether the SerS^wt^ protein could translocate into DeaD-marked condensates. As hypothesized, SHX treatment also induced the compartmentalization of SerRS^wt^ into DeaD-marked condensates in KLY cells, resembling the mutant *SerS*^*T*^ phenotype (Fig. 4f, 4g). Consistently, serine supplementation reversed this effect, releasing a substantial fraction of SerRS^T^ from DeaD-granules (Fig. 4f). Together, these findings demonstrate that serine starvation triggers sequestration of SerRS^T^ into DeaD-marked condensates, regulating dormancy and driving persistence.

## DeaD undergoes phase separation to sequestrate SerRS^T^

Since LLPS is a critical mechanism underlying biomolecular condensate formation in bacteria and eukaryotes (*33, 43*–*46*), we investigated whether DeaD undergoes liquid-liquid phase separation (LLPS) to form condensates, thereby selectively partitioning SerRS^T^ over SerRS^wt^. Fluorescence recovery after photobleaching (FRAP) measurements revealed that DeaD-condensates in the *SerS*^*T*^ strain showed significantly slower recovery kinetics and a higher immobile fraction, indicating that SerRS^T^ perturbs the dynamics of DeaD-condensates *in vivo* (Fig. 5a). This effect was particularly pronounced in M9 medium, coinciding with the prolonged lag phase observed in mutant cells (Fig. S6a). *In vitro*, DeaD protein underwent phase separation, dependent on protein and salt concentrations (Fig. 5b) (*38*). We then confirmed that DeaD condensates coalesce (Fig. 5c), and FRAP showed recovery of fluorescence signal (Fig. S6b), indicating their liquid-like material properties. We hypothesized that DeaD condensates function as scaffolds, whereas SerRS^T^ protein acts as a partitioning client molecule. Consistent with this, we observed that the purified SerRS^T^ protein was significantly enriched by partitioning into DeaD condensates (Fig. 5d, S5c-S5e), suggesting that LLPS-mediated recruitment drives the sequestration of SerRS^T^ into DeaD-condensates.

**Fig. 5.**
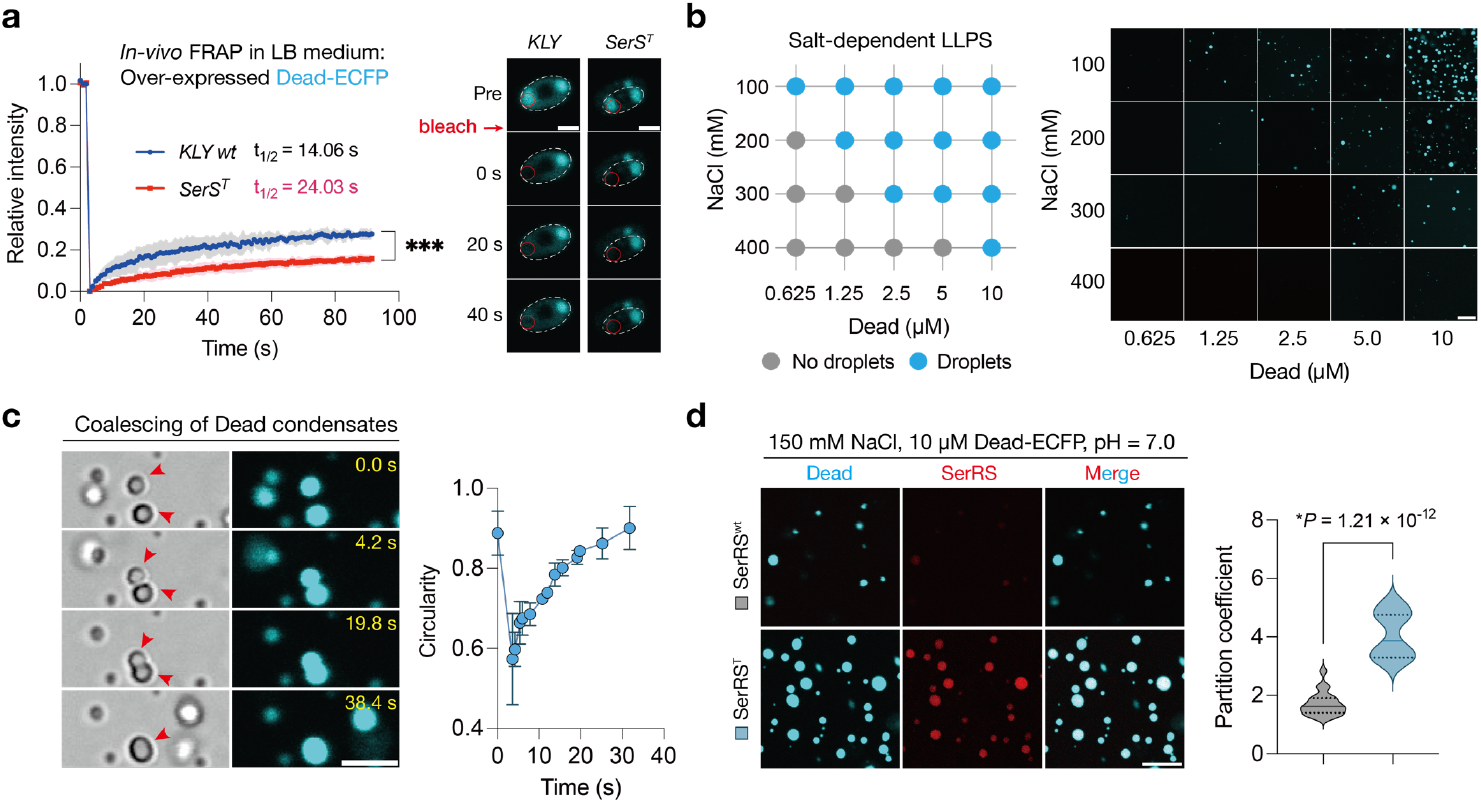
DeaD condensates selectively recruited the SerRS^T^ protein through LLPS. (a) FRAP analysis of DeaD-granules in the ancestral KLY and *SerS*^*T*^ mutant cells cultured in LB medium. Data are mean ± s.d., ***p < 0.001 by two-way ANOVA; Sidak’s multiple comparison test. Mobile fractions were 27% (KLY *wt*), 16% (*SerS*^*T*^). Scale bar, 1 μm. (b) LLPS of purified recombinant DeaD at various amounts and NaCl concentrations. Left panel, summary of phase-separation behavior of protein DeaD; right panel, representative fluorescent microscopy images. Scale bar, 20 μm. (c) Coalescing DeaD condensates. Left panel, time-lapse imaging. Right panel, circularity of fusing condensates over time. Data are mean ± s.d., n = 3 condensates. Scale bar, 5 μm. (d) SerRS^T^ partitioning by DeaD condensates *in vitro*. Left panel, microscopy images showing the enrichment of purified SerRS^T^ protein (red font and image, with a C-terminal mCherry tag) by partitioning to DeaD condensates (cyan font and image) *in vitro*. Right panel, quantification of 561 nm emission inside and outside of DeaD condensates. Solid lines indicate median. *P* values were calculated using two-tailed t-tests; n = 20 condensates, respectively. Scale bar, 10 μm.

### SerRS^T^ sequestration impairs localized translation in DeaD-condensates

Biomolecular granules are considered to regulate translation by compartmentalizing translation machinery or modulating mRNA dynamics, yet direct evidence for these functions remains insufficient (*36, 37, 47*). To investigate whether the perturbed Dead-granules impact translation, we employed surface sensing of translation (SUnSET), where puromycin incorporates into nascent polypeptides by mimicking an aminoacyl-tRNA (*48*), to monitor the global protein synthesis in the KLY and *SerS*^*T*^ strains. In *SerS*^*T*^ strain, translation patterns shifted significantly, accompanied by a negligible reduction in global translation efficiency compared to KLY (Fig. 6a, 6b). To assess the spatial distribution and intensity of *in situ* translation, we used o-propargyl-puromycin (OPP) coupled with click-reaction labeling to visualize OPP-labeled polypeptides (*48*). In ancestral cells, translation hotspots were clearly identifiable, whereas these sites were absent in *SerS*^*T*^ mutants (Fig. 6c). DeaD-condensates strongly colocalized with translation hotspots in ancestral cells, suggesting their role as hubs for localized protein synthesis (Fig. 6d, 6e, top). In contrast, active translation sites largely absent from DeaD-condensates in *SerS*^*T*^ cells (Fig. 6d, 6e, second row). Consistent with previous hypotheses, SHX treatment specifically impaired DeaD-localized translation activity in wild-type KLY cells, while serine supplementation restored translation hotspots in the *SerS*^*T*^ cells (Fig. 6d, 6e). These findings demonstrate that DeaD-condensates may serve as platforms for localized translation, with aberrant SerRS sequestration, triggered by SerRS^T^ or SHX exposure, impairing their function and contributing to dormancy and antibiotic persistence.

**Fig. 6.**
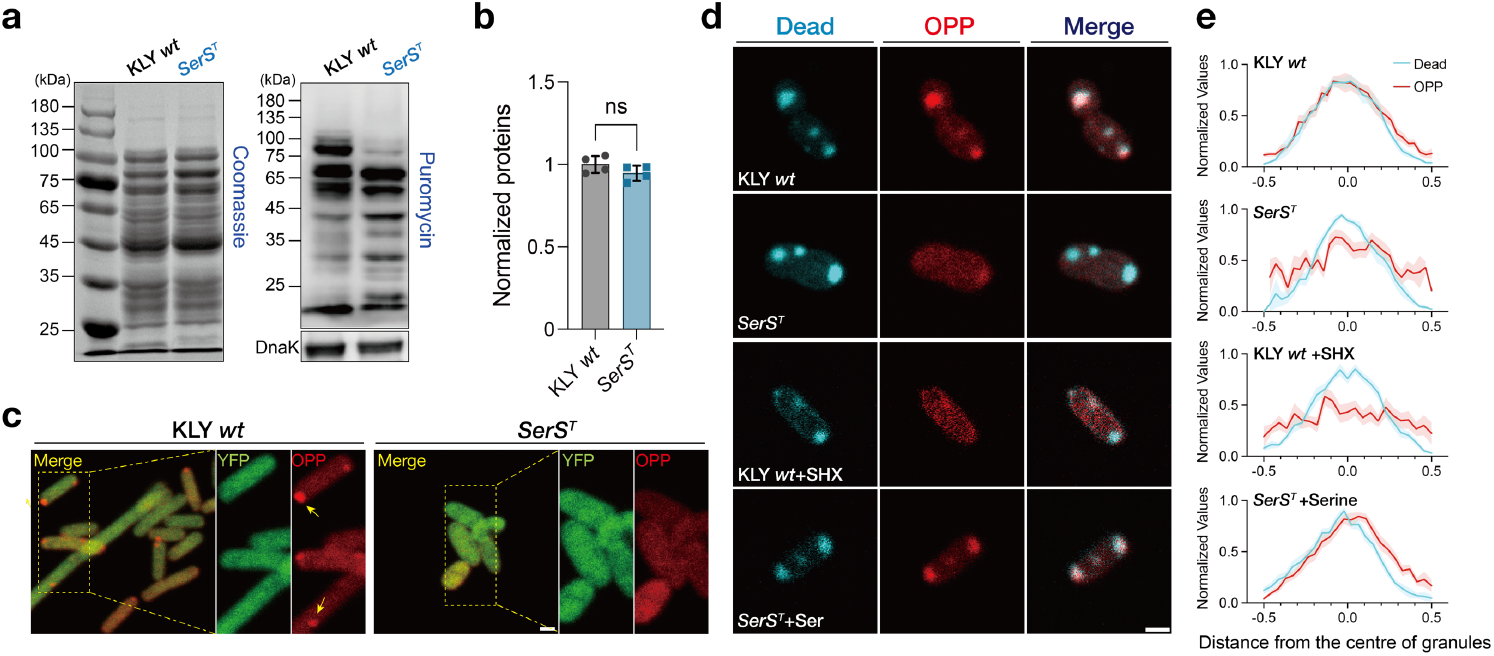
Aberrant recruitment of SerRS^T^ impaired localized translation in DeaD-condensates. (a) Western blot analysis showing newly synthesized polypeptides labeled by puromycin in KLY *wt* and *SerS*^*T*^ cells. Labeling time: 30 min. Total protein is shown as a reference for normalization. (b) Quantification of puromycin incorporation normalized to total protein. Individual data points represent biological replicates, data are represented as the mean ± s.d.; *P* value was calculated using an unpaired t-test. (c) Microscopic images of the KLY *wt* and *SerS*^*T*^ cells treated with OPP and stained with Alexa 594, illustrating active translation upon serine starvation. Scale bar, 1 μm. (d) Co-localization of nascent peptides and DeaD-condensates in KLY *wt* and *SerS*^*T*^ cells expressing ECFP-tagged DeaD. Altered co-localization patterns were observed in SHX-treated KLY *wt* cells and the serine-restored *SerS*^*T*^ cells. Scale bar, 1 μm. (e) Signal intensity profiles of OPP-labelled nascent polypeptides (red), centered at DeaD-condensates (cyan). Shaded areas around lines indicate SEMs (n = 10,10,11,13).

### SerRS^T^ disrupts translational composition by partitioning into DeaD-condensates

LLPS-driven biomolecular condensates, such as P-bodies and stress granules, dynamically alter their protein composition in response to environmental changes or genetic mutations (*36, 49*). To investigate how SerRS^T^ sequestration disrupts the translational function of DeaD-condensates, we performed proteomic analysis of the insoluble fraction from the KLY and *SerS*^*T*^ tolerant strains, identifying approximately 500 proteins with altered translocation (Fig. 7a). Gene functional classification revealed that many of these proteins are essential components of translational machinery, including aminoacyl-tRNA synthetases (aaRSs), ribonucleoprotein complexes, translation factors, and ATP-dependent helicases (Fig. 7a). These findings were further validated in the SerRS-associated interactome using co-immunoprecipitation, a method that preserves the integrity of protein complex (Fig. S7a-7c). Gene ontology (GO) analysis of SerRS-immunoprecipitated proteins and insoluble condensate fractions demonstrated strong enrichment of translational processes and ribosomal subunit assembly in the *SerS*^*T*^ strain relative to the ancestral KLY strain (Fig. 7b).

**Fig. 7.**
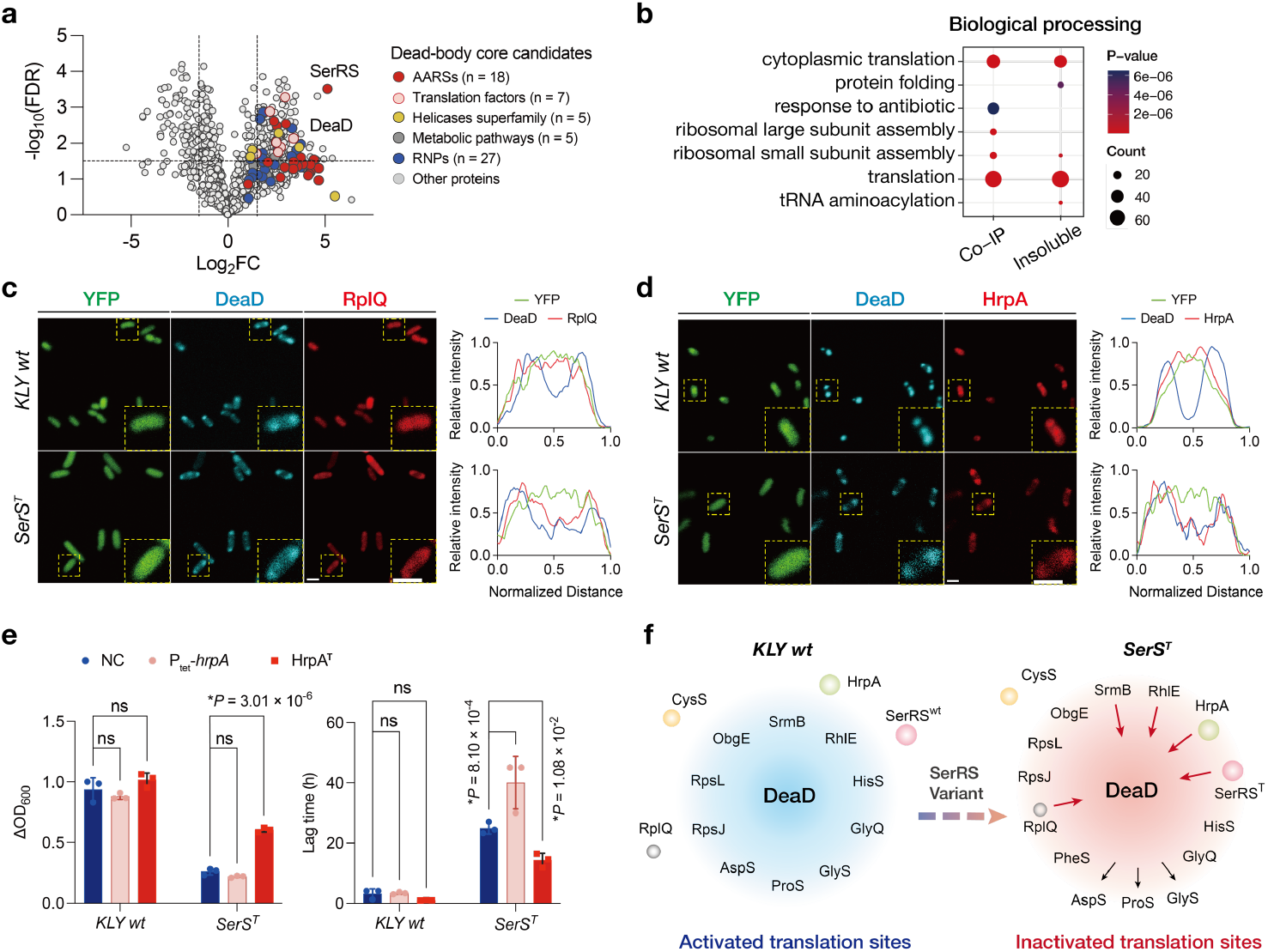
Perturbation of the translational network within DeaD-marked condensates due to SerRS^T^ sequestration. (a) Volcano plots displaying MS analysis of insoluble fractions from the *SerS*^*T*^ strain relative to KLY *wt* strain. Statistical significance was assessed using a t-test with the false discovery rate (FDR) correction adjusted p < 0.02. (b) Gene ontology analysis of immunoprecipitated proteins from the SerRS^T^ versus SerRS^wt^ interactomes, and components enriched in the insoluble pellets of mutated versus wild-type cells. (c-d) Representative images of mCherry-tagged ribonucleoprotein RplQ (c) or HrpA (d) in ancestral and *SerS*^*T*^ cells expressing ECFP-tagged DeaD. Inset: higher magnification of the yellow boxed area. Scale bar, 1μm. Line scans show the intensity profiles of DeaD co-localized with mCherry-tagged candidates. (e) Effect of HrpA mutants on the growth defect. Expression plasmid of HrpA was induced by 0.05 μg/mL tetracycline and the truncated mutation of *hrpA* was generated in the genomes of both *SerS*^*T*^ and ancestral strains via the CRISPR-Cas9 system. Data are mean ± s.d.; *P* values were calculated using two-tailed t-tests. (f) Schematic depicting the altered translocation of translational components within DeaD-marked condensate. See also Fig. S8.

We hypothesized that the translocated candidates, whose expression levels critically impact cell growth in the mutant strain, may result from the abnormal recruitment of SerRS^T^ protein. Several candidate proteins were selected to assess their effects on the growth phenotype of the *SerS*^*T*^ strain. Notably, overexpression of certain ribosomal subunit, particularly L17 RplQ (encoded by *rplQ*), significantly rescued the growth defect in the *SerS*^*T*^ strain (Fig. S7d). Fluorescence microscopy further revealed enhanced co-localization of RplQ with DeaD-marked condensates in *SerS*^*T*^ cells compared to wild-type cells (Fig. 7c, S7f). To identify genes contributing to the SerRS^T^-driven phenotype, an inverse selection strategy was employed via cyclic batch culturing to enrich mutants with a growth advantage over the *SerS*^*T*^ strain (Fig. S7e). In addition to *serS*^*T*^, these growth-restored mutants could harbor a second mutation that potentially counteract with SerRS^T^ function. Among these, a truncated version of HrpA (designated HrpA^T^, c.2684delA) was identified, whose wild-type allele was also enriched in the SerRS^T^ interactome (Fig. S7c). *E. coli* HrpA, an ATP-dependent RNA helicase of the DEAH/RHA family, has been implicated in modulating antibiotic susceptibility, possibly through effects on RNA dynamics and bacterial stress responses (*50*–*52*). Fluorescence imaging showed that mCherry-tagged HrpA colocalized with DeaD-marked condensates in *SerS*^*T*^ cells, whereas it displayed a diffuse cytoplasmic distribution in wild-type cells (Fig. 7d, S7g). Moreover, overexpression of HrpA exacerbated the growth defect in the *SerS*^*T*^ strain, a phenotype that was partially restored in the *SerS*^*T*^*hrpA*^*T*^ mutant (Fig. 7e). Additionally, DeaD interactions with representative aaRSs in the mutant and ancestral cells were also confirmed. HisS, GlyQ, PheS represented constant or increased interactions with DeaD-condensates in mutant cells, while GlyS, AspS and ProS as representatives exhibited reduced interactions (Fig. 7f and Fig. S8). Notably, CysS was consistently excluded from DeaD-driven granules. These findings suggested that SerRS^T^ sequestration modulated translational components within DeaD-marked condensates, including RplQ, HrpA and various aaRSs, thereby impairing their function and contributing to dormancy and persistence.

## Discussion

We tracked the evolution of exponential-phase populations under cyclic antibiotic treatment and identified a serine starvation-triggered persistence phenotype mediated by *serS* mutations, akin to the response seen in SHX-treated cells. Exhaustion of serine induced the aberrant translocation of SerRS into DeaD-marked condensates, perturbing the interactome and impairing localized translation hotspots. Both the *SerS*^*T*^ mutant and SHX-treated KLY strains exhibited similar patterns of SerRS sequestration, accompanied by a substantial extension of lag time upon transfer to amino acid-free media. In natural environments, as nutrient availability fluctuates between periods of feast and famine, bacteria likely integrate various environmental cues to establish the transition between growth and dormancy, such as the duration of dormancy and the timing of division. The phenomenon of sharp serine consumption during the exponential phase has been well-documented for decades (*18*) and may serve as an indicator of nutrient consumption, allowing bacteria to adjust their population size. In eukaryotes, aaRSs can form multi-synthetase complexes (MSCs), which are increasingly recognized as mediators of cell signaling for extra-translational functions (*53, 54*). Since the identification of high-persistence mutants, such as HipA7 and *MetG*^*T*^ (*13, 14, 28*), how the length of lag time during dormancy is regulated by biological pathways involving aaRSs remains inadequately understood. Our results suggest a potential common mechanism, wherein aaRSs sense the concentrations of cognate amino acids and partition in biological condensates formed through LLPS, thereby regulating cell fate. We propose the term “Translation-localized Dormancy Granule” to describe these structures.

LLPS-mediated biomolecular condensates are membrane-less intracellular assemblies that are tightly regulated in response to environmental signals. In eukaryotes, DEAD-box ATPases (DDXs) have emerged as conserved and redundant regulators of RNA-containing condensates that mediate stress response and control cell fate (*37, 39, 49, 55, 56*). Our findings suggest that these DeaD-marked granules evolved in much simpler organisms than previously appreciated, enabling adaptive growth transitions in response to nutrient fluctuations. We were particularly struck by the observation that these condensates are involved in localized translation activation, consistent with similar findings in eukaryotic cells (*37, 47, 57*). Furthermore, translational factors identified within DeaD-marked condensates here are conserved components of most known granules (*35, 41, 58*). RNA post-transcriptional processing is another plausible function of these condensates. Although RNA splicing is rare in prokaryotes, RNA editing, degradation, and tRNA modification are essential for gene regulation (*41, 59*). For example, HrpA, which processes the *daa* operon mRNA in *E. coli*, shares significant sequence similarity with yeast RNA helicases (PRP2, PRP16, and PRP22) involved in mRNA splicing (*50, 51, 60, 61*). These associations imply that these condensates might have evolved in prokaryotes to selectively modulate translation and combat various stresses.

Cell-to-cell heterogeneity is generally attributed to intrinsic gene expression noise (*62, 63*), in which condensate formation has been suggested as a potential mechanism to reduce cytoplasmic noise (*64*), or conversely, to amplify cell-cell variability of condensate regulated cellular physiology properties (*65*). SHX-induced perturbations were predicted to disrupt the cellular network, shifting its dynamics from biological adaptation toward the generic behavior of a large random network, rather than a specific genetic pathway (*22*). Inspired by this hypothesis, the analogy of serine starvation-triggered persistence mediated by SerRS^T^ mutation and SHX treatment led us to explore a similarly dysregulated network, which could explain the extensive heterogeneity observed in the dormancy duration of triggered persistence.

The identification of condensate-regulated localized translation in prokaryotes suggests that LLPS-driven, membrane-less granules may represent an evolutionarily conserved phenomenon, providing a versatile model for studying the mechanisms and evolution of stress responses and cell fate decisions. This insight could also pave the way for new strategies targeting dormancy regulation and therapeutic intervention in persistence-associated infections.

## Acknowledgement

We thank Jingjing Wang at the Cell Biology Facility, Center of Biomedical Analysis, Tsinghua University, for her assistance with confocal microscopy. We also thank Engineer Weihua Wang from Center of Pharmaceutical Technology, Tsinghua University, for quantifying amino acids concentrations using LC-MS. Our gratitude extends to the Tsinghua University Technology Center for Protein Research for their mass spectrometry services and MS data analysis. The CRISPR/Cas9-based genome editing tool were generously donated by Huo’s lab, School of Life Science, Beijing Institute of Technology. We appreciate Nathalie Balaban’s lab at Racah Institute of Physics, The Hebrew University of Jerusalem, for providing the *E. coli* KLY strain. We also thank Pilong Li from the School of Life Sciences, Tsinghua University, for his guidance on phase separation assays. Additionally, we are grateful to Xiaofeng Fang (School of Life Sciences, Tsinghua University) and Junhao Zhu (Institute of Microbiology, Chinese Academy of Sciences) for their suggestions. This research was supported by the National Key Research and Development Program of China (2022YFC2303202), the Tsinghua-Peking Joint Center for Life Sciences (20111770319), and the Tsinghua University Dushi Program (20221080037).

## Author Contributions

J.L. mainly designed experiments, acquired funding and revised the paper. Z. Z. designed and performed experiments, analyzed data and wrote the paper. D. L. performed the *in-vitro* phase separation assay. B. Z. analyzed the whole-genome sequencing results.

## Supplementary information

Supplemental Information contains eight figures, two tables and two videos.

