## Supplementary material for "Sequestration of SerRS through LLPS Impairs Localized Translation and Contributes to Antibiotic Persistence": SupplementalFiles.docx

**The PDF file includes:**

Methods

Fig. S1 to S8

Tables S1 to S2

**Other Supplementary Material for this manuscript includes the following:**

Supplemental Movies S1-S2

**Methods**

**Strains and cell culture**

The ancestral strain (KLY) was conducted by P1 transduction of the yfp-Cam cassette form M22 into *E. coli* KL16, as previously described (*1*)**.** All experiments were conducted at 37°C in LB (Sigma) medium, shaking at 220 rpm, unless otherwise specified. Bacterial cultures were grown to exponential phase and stored in aliquots with 15% glycerol at -80°C. Cell growth in M9 medium (Sangon, M9 minimal salts) was initiated by first growing the cultures in LB medium until ∆OD_600_ ≈ 0.1 (following serine starvation), after which they were transferred to M9 medium via a 1:1000 dilution. The cultures were then incubated at 37°C with shaking at 220 rpm.

**Construct**

The expression plasmids of SerRS^wt^-mCherry and SerRS^T^-mCherry were constructed as follows. DNA fragments encoding SerRS^wt^ and SerRS^T^ were PCR-amplified from the KLY *wt* and *SerS^T^* genomes, respectively, and inserted into a p15A-Ptet-Kan^R^ backbone using the ClonExpressⓇII One Step Cloning Kit (Vazyme, C112-01). To generate DeaD expression plasmids, the DNA fragment encoding DeaD was PCR-amplified from the KLY *wt* genome, fused to an ECFP tag, and assembled into a pET28a backbone (Amp^R^), with expression driven by a *P_lac1_* promoter induced by Isopropyl β-D-thiogalactopyranoside (IPTG). His-tagged DeaD was assembled into a p15A-Ptet-Kan^R^ backbone. Other DEAD RNA-dependent ATPases, including SrmB, RhlE and HrpA, were cloned from the KLY genome and assembled into the p15A-Ptet-Kan^R^ backbone. DNA fragments encoding RNPs were PCR-amplified from the KLY *wt* genome, fused to mCherry, and inserted into a pET28a backbone (Kan^R^). The aaRS genes (GlyQ, GlyS, CysS, PheS, ProS, AspS, HisS) were cloned from the KLY *wt* genome, fused to mCherry, and the assembled into the p15A-Ptet-Kan^R^ backbone. All constructs were confirmed by sequencing.

**Cyclic Evolution Design**

The cyclic experimental design comprised three main steps. First, 50 μL aliquots (approximately 10^7 CFU/mL) was resuspended into 50 mL LB and incubated until the optical density (OD_600_) reached approximately 0.12. Second, the cultures were supplemented with 2 μg/mL ertapenem (about 100-fold MIC) (Shanghai yuanye Bio-Technology Co., Ltd) for two hours. Finally, the antibiotic-containing supernatant was removed by centrifugation (6000 rpm for 6 minutes), and the cell pellet was washed twice with 1 mL of 0.9% NaCl. The cell pellets were resuspended in 50 mL fresh LB and incubated until they reached exponential phase (OD_600_ ≈ 0.12). MIC values was determined using serial 2-fold dilutions, with shaking for 24 h at 37 °C. MIC was defined as the lowest concentration of antibiotic that inhibited bacterial growth.

**Sampling**

Unless otherwise specified, LB-cultured cells used in this study were collected at an ∆OD_600_ of approximately 0.1 (OD_600_ ≈ 0.2) following serine starvation. At this point, the *SerS^T^* tolerant strain exhibited an abrupt growth arrest, while the KLY strain continued to divide.

**ScanLag**

A sample of the culture was serially diluted and plated onto solid LB agar. These plates were then incubated in a ScanLag system (*2*) at 37°C, an array of office scanners that automatically captures images every 20 minutes to monitor the appearance of thousands of colonies. An automated image analysis tool processes the images to derive the distribution of colony appearance times.

**Killing assay**

Bacteria cultures were grown in LB to OD_600_ ≈ 0.12, supplemented with 2 μg/mL ertapenem and incubated for three hours at 37°C shaking at 250 rpm. CFUs were determined by plating before and after antibiotic treatment. All assays were performed in triplicate.

**Whole genome sequencing**

Strains were grown overnight in LB medium, and genomic DNA was extracted using the TIANamp Bacteria DNA kit (DP302). DNA samples from each strain was sent to Beijing Saimo Baihe Biotechnology Co., Ltd. for Illumina sequencing. Genomic analysis was performed using Geneious Prime to detect SNPs and new junctions. Each reported SNP was confirmed by PCR amplification and sequencing.

**Confocal microscopy**

Time-lapse microscope for recording bacterial growth	Cells in an exponential phase were collected, washed three time with LB medium and imaged on a gel pad containing 1% agarose. The LB medium gel-pad was prepared as a gel island at the center of a confocal dish. Cells were monitored under DIC (differential interference contrast, Nikon Ti2-E) microscopy at 37°C using a live-cell system. To observe bacteria resuscitation under the minimal M9 gel-pad, LB-cultured cells were collected, washed three times with M9 minimal medium and placed under a fresh M9 gel-pad as described above. The cells were then imaged under bright-field and fluorescence illumination (Nikon AXR NSPARC) at 37°C.

Microscope imaging	Images were captured using confocal imaging (Nikon AXR NSPARC) with a 100× oil objective. ECFP was excited at 445 nm, YFP at 488 nm, mCherry/ Alexa 594 at 561 nm, Alexa 594 at 594 nm. The fluorescence signals were collected using filter sets: 474/20 nm for ECFP, 540/20 nm for YFP, 618/20 nm for mCherry/ Alexa 594. Colocalization analysis was performed using Fiji (Colocalization Finder). A square region containing KLY *wt* or *SerS^T^* cells with condensates was selected as the region of interest (ROI). Pearson’s correlation coefficient between two channels within the ROI was calculated to quantify colocalization. For line scan analysis, a straight line was drawn across the enlarged image the ROI and the intensity profile was measured using the Plot Profile tool in Fiji.

Translation visualization	Cells were incubated with O-propargyl-puromycin (OPP) using Click-iT™ Plus OPP Alexa Fluor™ 594 Protein Synthesis Assay Kit (Invitrogen) (*3*) according to the manufacturer’s guidelines. Briefly, cells we incubated with 20 μM OPP for 30 min at 37 °C with shaking. Cells were then fixed with 4.0% formaldehyde and permeabilized with 0.1% Triton X-100 for 15 min at room temperature. Fluorescent labelling was performed for 30 min with Alexa Fluor 594 reaction cocktail at room temperature.

**Detecting amino acids concentrations**

Extracellular and intracellular amino acids concentrations were quantified by liquid chromatography and mass spectrometry (LC-MS). Metabolites were sampled at regular intervals during the testing phase and extracted using 80% methanol. The extracts were directly analyzed without drying. Metabolite quantification was performed using a 4500+QTrap mass spectrometer (AB SCIEX, USA) and an ACQUITY UPLC H-Class system (Waters, USA). Chromatographic separation was achieved using an ACQUITY UPLC BEH Amide column (2.1×100 mm, 1.7 μm; Waters). Mobile phase A contains HPLC-grade ACN and 0.1% formic acid, while mobile phase B is water and 0.1% formic acid. Data were collected in positive mode using the Multiple Reaction Monitoring (MRM) mode with an ESI source. The gas pressure for the atomizer (gas 1), heater (gas 2) and curtain were set to 55, 55 and 30 psi respectively. The ion spray voltage for the positive ion mode is 5,500 volts. The optimal probe temperature was determined to be 500°C, and the column oven temperature was set at 40°C.The AB SCIEX 1.6.3 analyzer (AB SCIEX, USA) was used for metabolite identification and peak integration.

**Insoluble protein isolation and SDS-PAGE**

Insoluble proteins were isolated from bacterial cultures following a modified method from Tomoyasu and Pu (*4*, *5*)**.** Briefly, 20 mL of bacterial culture (OD_600_ = 0.2) was rapidly cooled on ice. Cells were collected by centrifugation at 6,000 g for 10 minutes at 4°C. The pellet was resuspended in 40 μL buffer A (10 mM potassium phosphate buffer, pH 6.5, 1 mM EDTA, 20% w/v sucrose, 1 mg/mL lysozyme) and incubated on ice for 30 minutes. The cell lysate was mixed with 360 μL buffer B (10 mM potassium phosphate buffer, pH 6.5, 1 mM EDTA) and subjected to sonication on ice. Membrane proteins were dissolved by resuspending the pellet in 400 μL of buffer C (buffer B with 2% NP40). The insoluble proteins were collected by centrifugation at 15,000 g for 30 minutes at 4°C. NP40-insoluble pellets were washed twice with 400 μL of buffer B and resuspended in 50 μL buffer B by brief sonication. Proteins were separated by 10% SDS-PAGE and stained with Coomassie Brilliant Blue.

**Western Blots**

Protein concentrations were determined using a BCA protein assay kit (Beyotime, P0012). Proteins were separated on SDS-PAGE and transferred to a PVDF membrane. The membranes were blocked in PBS containing 0.5% milk and 0.5% Tween20 for 1 hour at room temperature. Primary antibodies were incubated overnight in PBST containing 4% BSA, 1% Tween20, 0.05% NaN3, then incubated with the appropriate HRP-conjugated secondary goat antibodies (Beyotime, A0350) for 2 hours at room temperature, following by extensive washing. Blots were detected using GE western blotting substrate, and acquired images were analyzed with ImageJ software. Primary antibodies used in this study were mCherry (Cell Signaling, 43590), His-tag antibody (Sangon, D191001), Anti-DnaK antibody (Abcam, ab80161), anti-puromycin (AbClonal, A23031).

**Co-immunoprecipitation**

Co-immunoprecipitation was performed using KLY wild-type and *SerS^T^* strains harboring an mCherry tag fused to the C-terminus of SerRS protein. Cultures were grown to OD_600_ ≈ 0.2 (upon serine depletion) and lysed using lysis buffer (50 mM NaCl, 20 mM Tris-HCl, pH 7.4) containing a protease inhibitor cocktail (Beyotime, P1005), 1 U/mL Superase IN (Invitrogen, AM2694), and 10 U/mL RNase-free DNase I (Thermo, EN0521). Immunoprecipitation was performed with Anti-mCherry Nanobody Immunomagnetic Beads (Elabscience, EAIP-005MN). The precipitates were subjected to the quantitative mass spectrometry to analyze the interacted protein candidates.

**Mass spectrometry**

The SDS-PAGE gel was reduced with 5 mM of dithiothreitol and alkylated with 11 mM iodoacetamide, followed by in-gel digestion with sequencing grade modified trypsin at 37°C overnight. Peptides were extracted twice with 0.1% trifluoroacetic acid in 50 % acetonitrile aqueous solution for 30 min and centrifuged in a SpeedVac to concentrate the extracts. The digestion products were separated via a 120-min gradient elution at a flow rate 0.300 µL/min on a Thermo Scientific UltiMate™ 3000 HPLC system, directly interfaced with the Thermo Orbitrap Fusion Tribrid mass spectrometer. The analytical column was a C18 fritless column (75 μm x15cm, Acclaim PepMap nanoViper RSLC Thermo). Mobile phase A consisted of 0.1% formic acid, and mobile phase B consisted of 80% acetonitrile and 0.1% formic acid. The Orbitrap Fusion was operated in data-dependent acquisition mode using Xcalibur 4.1 software, performing a full-scan mass spectrum (350-1550 m/z, 120,000 resolution) followed by top-speed MS/MS scans in the Orbitrap. The MS/MS spectra were searched against *Escherichia coli* K-12 database using Proteome Discoverer (Version PD3.1, Thermo-Fisher Scientific, USA).

**CRISPR-Cas9 genome editing**

Genome editing of *serS^wt/T^*, *dead*, and *hrpA* was performed using the CRISPR/Cas9 system(*6*). Briefly, the plasmid Kan^R^ p15A-PBAD-Cas9-PT5-Red𝛄𝛽𝛂 (plasmid#1) was respectively transferred into the *E. coli* KLY *wt* and *SerS^T^* strains to obtained the corresponding transformants. Temperature-sensitive Amp^R^ plasmids were constructed to express specific sgRNA and generate specific donor DNA, these plasmids were collectively named pSC101-PBAD-sgRNA-Donor (Plasmid #2). The Plasmid #2 was introduced into competent cells (containing pHCY-25A) and resuscitated for 45 min. Then, cells were plated on LB agar containing kanamycin and ampicillin and cultured overnight at 30°C. Following primary selection, cultures were inoculated into LB broth (containing kanamycin and ampicillin) and cultured at 30°C for 2 hours. IPTG was added, and the mixture was cultured for 1 hour at 30°C; L-arabinose was then added and incubation continued at 30°C for 3 hours. The cultures were diluted 1:1000 into fresh LB, and cells were plated on LB agar plates containing kanamycin, ampicillin, with L-arabinose supplement. Colonies were selected for PCR validation. Plasmids were subsequently eliminated by culturing the bacteria on LB plates containing sucrose at 37°C. Knockout mutants were confirmed by colony PCR with optimized primers.

**Fluorescence Recovery After Photobleaching (FRAP)**

FRAP experiments were performed using an Olympus FV3000 confocal microscope. Condensates were bleached with a 405 nm Ar-laser pulse at maximum intensity. Data were analyzed using CellSens imaging software. Regions of interest (ROIs) were defined in the photobleached region, a non-photobleached cell, and the background. The mean intensity of each was extracted with the correction of photobleach and background, and the fit FRAP curves were generated.

***In vitro* protein expression and purification**

Fusion proteins were expressed in *E. coli* BL21 and purified using a Ni-NTA column. Protein expression was induced overnight at 16 °C with 0.5 mM IPTG. Cells were collected by centrifugation and resuspended in lysis buffer (20 mM Tris-HCl pH 7.4, 500 mM NaCl, 10% glycerol). The suspension was sonicated for 30 min (on 2 s, off 4 s, SCIENTZ) and centrifuged at 13,000g for 30 minutes at 4 °C. The supernatant was incubated with Ni-NTA agarose for 20 min, washed and eluted with 20 mM Tris-HCl pH 7.4, 500 mM NaCl and 500 mM imidazole. Eluted proteins were stored in 20 mM Tris-HCl pH 7.4, 500 mM NaCl, 1 mM dithiothreitol on ice or at −80°C.

***In vitro* phase-separation assay**

Purified DeaD protein was assembled by diluting the protein from a high-salt- storage buffer to varying salt concentrations (100-500 mM NaCl) and protein concentrations in 384-well plates (PhenoPlate™). DeaD-ECFP protein and SerRS^wt/T^-mCherry were gently mixed. After a 5-minute incubation, images were captured with an Olympus FV3000 confocal microscope equipped with a ×40 objective.

To quantify the **partition coefficient** of SerRS^wt/T^ proteins by DeaD condensates, regions of interest of equivalent size were analyzed in the Fiji implementation of ImageJ to calculate the SerRS signal intensity inside and outside DeaD condensates. The partition coefficient was defined as intra-condensate fluorescence intensity divided by the fluorescence intensity of the extra-condensate solution.

**Statistical analysis and data visualization**

Sample sizes are selected as widely used in the field. Biological and technical replicates were performed as described in the Methods for each experiment and conform to standards in the field. Two-tailed unpaired t-tests (with Welch’s correction, where applicable), assuming a parametric distribution, were performed to assess differences across at least three replicates. p<0.05 was considered significant. Statistical parameters including the definitions, exact values of n, and *P* values can be found in the figure legends and Methods. Data analysis was conducted using GraphPad Prism 10.2.0 Software, and final figures were assembled with Adobe Illustrator.

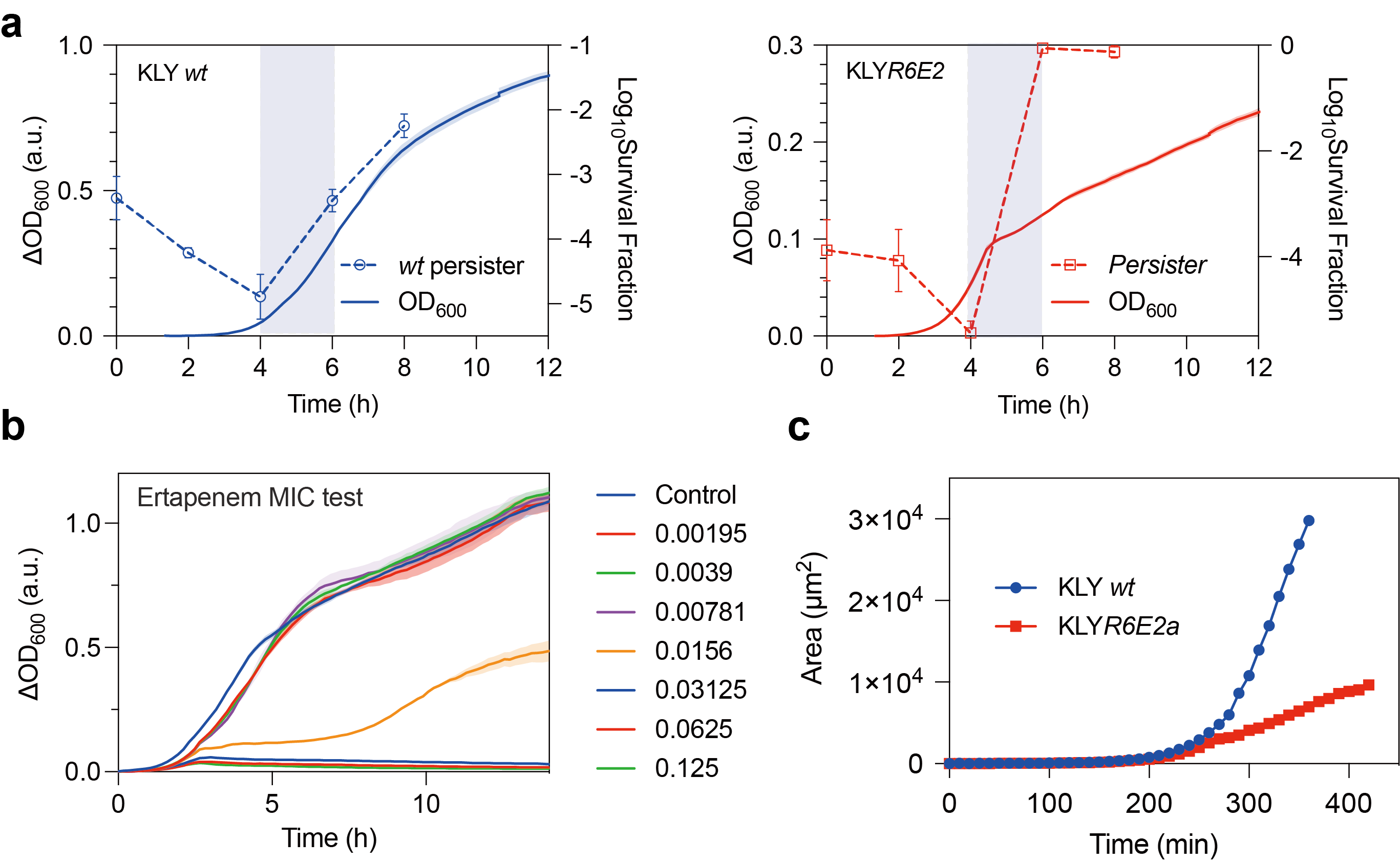

**Fig. S1 | A persistence mediated by growth arrest, related to Fig. 1**

(a) Total population and persisters cell count for the ancestor (left) and *SerS^T^* tolerant (right) strains over the course of cell growth. Cell cultures were sampled at an interval of 2 hours to conduct the antibiotic killing assay. (b) MIC of ertapenem determined via microdilution. The MIC of *E coli KLY* ranged from 0.0156 to 0.03125 μg/mL. (c) Growth area measured over incubation time on LB agar pads, observed under a microscope, corresponding to Fig. 1f.

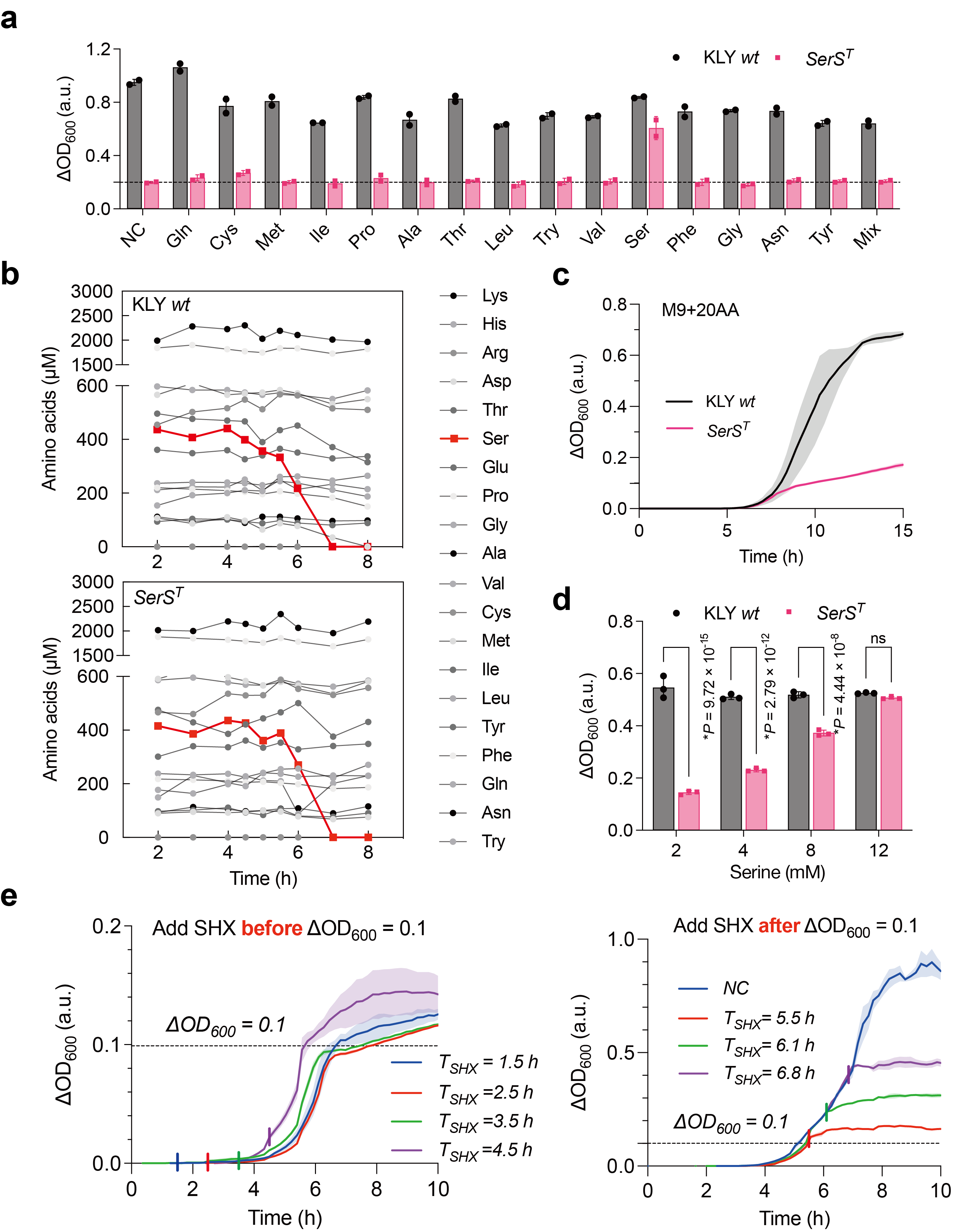

**Fig. S2 |** **Serine starvation triggered persistence, related to Fig. 2**

(a) Effect of various amino acid (AA) additions on the growth defect of the mutant strain in LB medium. Each amino acid was supplemented to a final concentration of 2 mM and finally cell densities were recorded after overnight incubation (approximately 16 hours). (b) Profiles of extracellular amino acid consumption by the ancestral and *SerS^T^* strains in LB medium. Data are derived from two biological replicates. (c) A similar growth defect of the *SerS^T^* strains displayed in defined M9 medium augmented with 20AAs, compared with that observed in LB medium. Each amino acid was added at a concentration of 2 mM. (d) Growth of the ancestral and *SerS^T^* strains in modified 20AAs-supplemented M9 medium with varying concentrations of serine added. (e) OD measurement of bacterial cultures exposed SHX during growth in LB medium. Left panel, SHX was introduced respectively at T=1.5, 2.5, 3.5, 4.5 hours, prior to serine deprivation, ∆OD_600_ < 0.1; Right panel, SHX was added after serine depletion, ∆OD_600_ ≥ 0.1. The colored solid lines indicate the addition of 1 mM SHX.

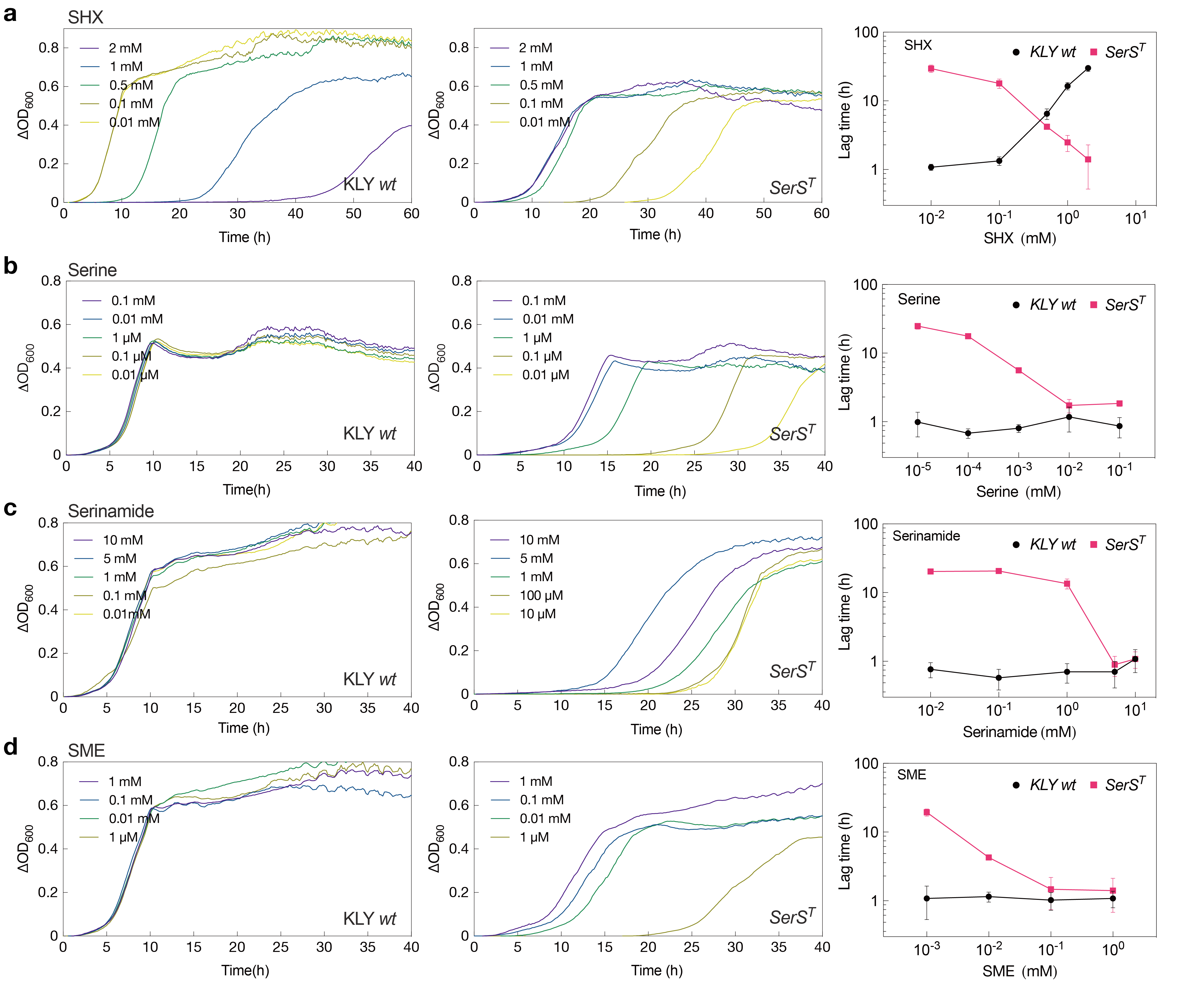

**Fig. S3 |** **Growth curves of the ancestor and *SerS^T^* strains upon transfer into M9 medium, with the exposure of serine and its analogues by varying levels of concentrations.** The lag time of bacterial cultures was calculated at the point when the ∆OD_600_ was less than 0.001, indicating minimal or no growth at the time of measurement. Data are represented as mean ± s.d. of three biological replicates.

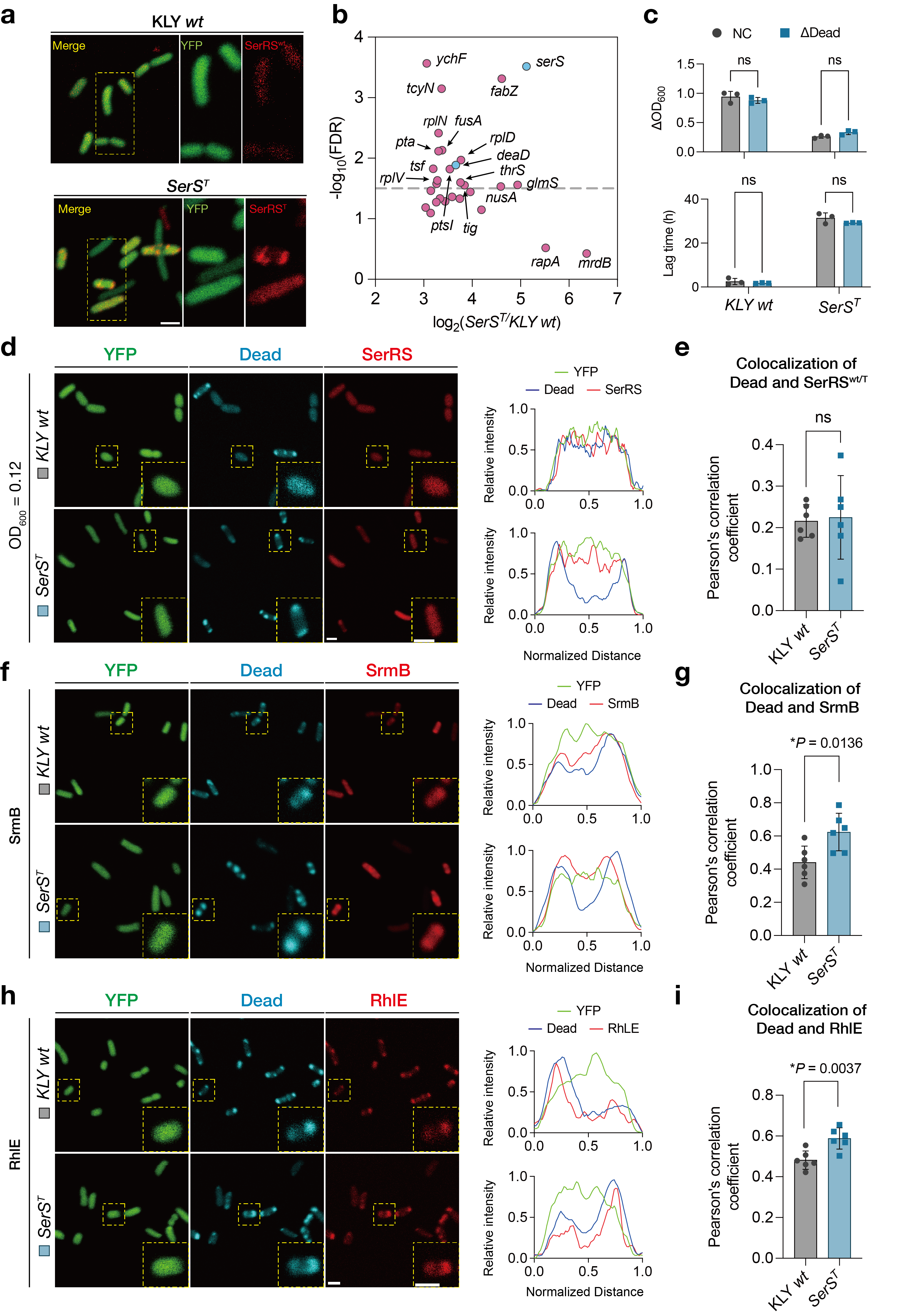

**Fig. S4 | SerRS^T^ aberrantly translocated into the DeaD-condensates upon serine deprivation, related to Fig. 4** (a) Fluorescence images of mCherry-tagged SerRS protein from *KLY wt* and the *SerS^T^* cells, with an enlargement of yellow framed region on the right. Scale bar, 2 μm. (b) Candidates highly enriched in the soluble pellet fractions of the *SerS^T^* strain relative to the ancestral KLY strain. Selection criteria: log_2_ *SerS^T^* (protein abundance of insoluble condensates in the mutant)>8, log_2_ (*SerS^T^*/KLY *wt*) (fold changes of the *SerS^T^* strain over KLY strain) > 3., as indicated in the upper-right corner of Fig. 4b. (c) Cell density in LB medium and lag time in M9 medium were measured in the ancestral and *SerS^T^* tolerant strains, both carrying a full-length knockout of the *deaD* gene. (d) Representative images of mCherry-tagged SerRS in cells expressing ECFP-tagged Dead before serine deprivation, at an ∆OD_600_ of approximately 0.01. Inset: higher magnification of the yellow boxed area. Scale bar, 2 μm. Line scans show the related intensity profiles of SerRS with DeaD, along with Pearson’s correlation coefficient (e). (f, h) The colocalization of other DEAD-box ATPases, SrmB and RhlE, with DeaD-marked condensates in the ancestor and *SerS^T^* mutant strains. (g, i) Pearson’s correlation coefficient showing the colocalization of DeaD-SrmB (g) and DeaD-RhlE (i). Data are presented as the mean ± s.d. with a two-tailed t-test.

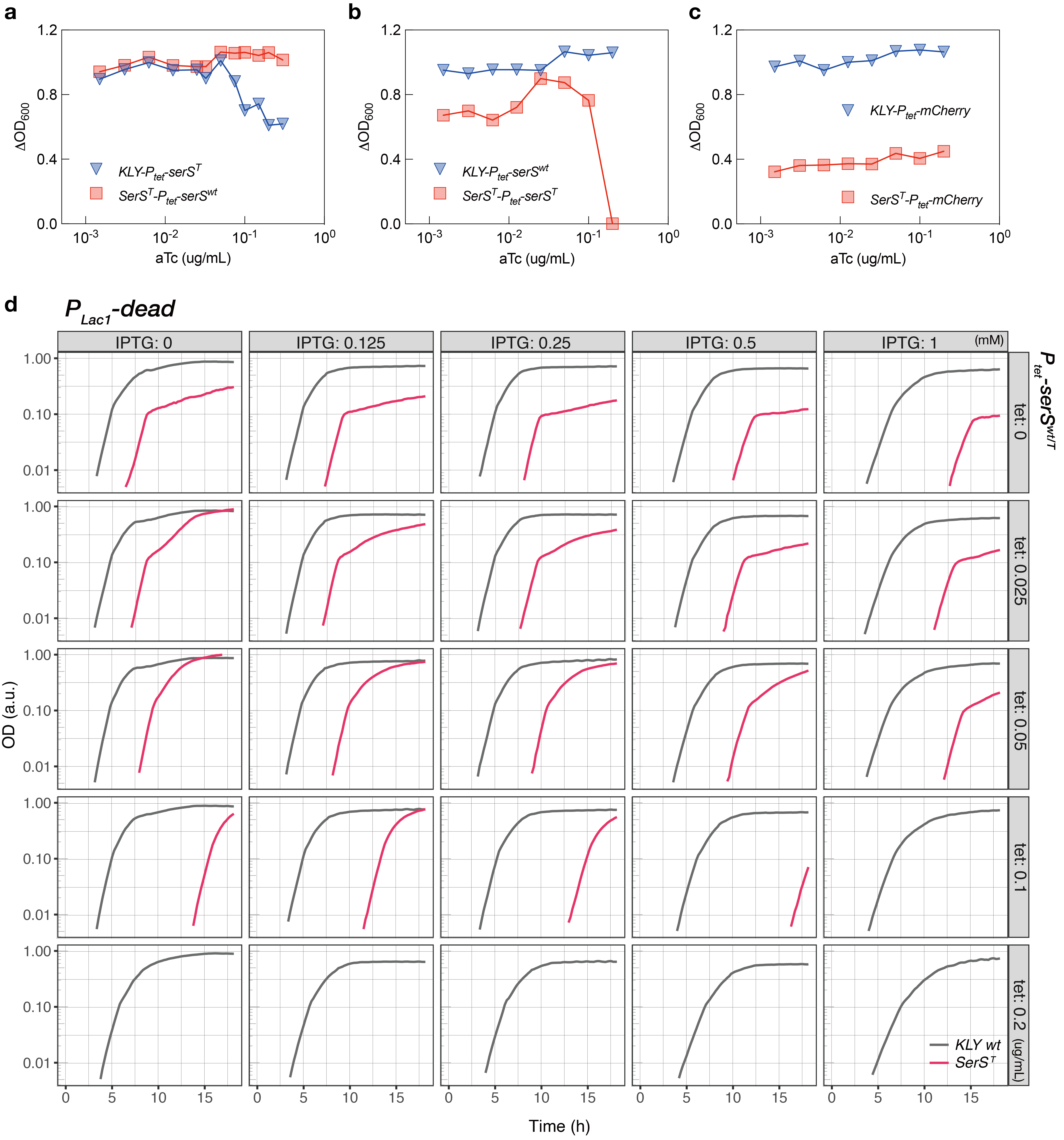

**Fig. S5 |** **DeaD and SerRS^T^ function as regulators of growth dynamics with opposing effects**

(a) Cell growth recovery in the *SerS^T^* tolerant strain by different levels of tetracycline. (b) OD measurement over incubation time with the different expression levels of DeaD and SerRS in the ancestral KLY and *SerS^T^* strains in LB medium. The KLY *wt* and *SerS^T^* strain co-expressed the DeaD and the respective *serS^wt^* or *serS^T^* gene, induced by varying levels of IPTG and tetracycline.

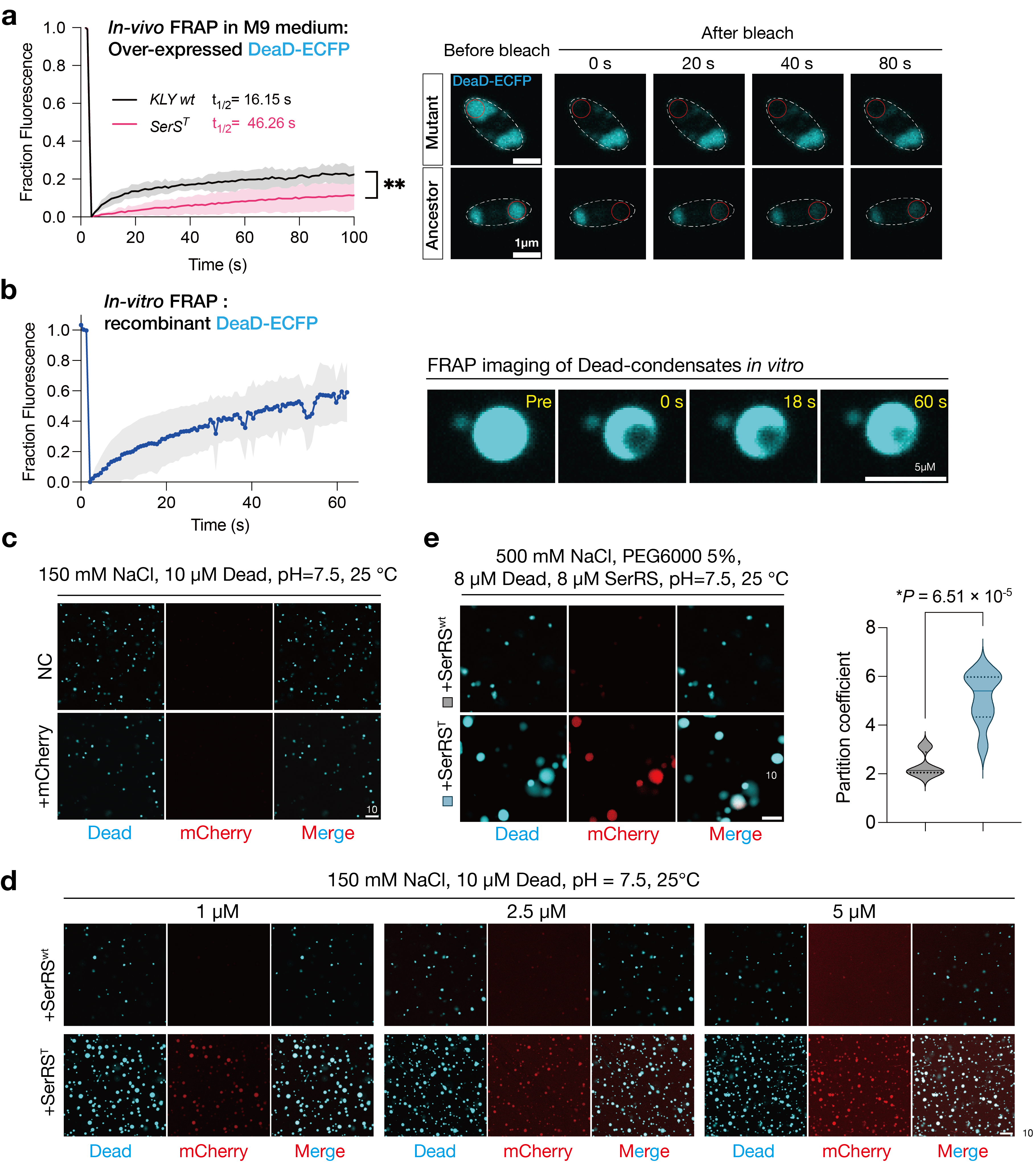

**Fig. S6 | DeaD undergoes phase separation to sequestrate SerRS^T^**

(a) FRAP analysis of DeaD-granules in the ancestral KLY and *SerS^T^* mutant cells cultured in M9 medium. Data are presented as mean ± s.d.; **p < 0.01 by two-way ANOVA; Sidak’s multiple comparison test. Mobile fractions were quantified at 25% (KLY *wt*), 18% (*SerS^T^*). Scale bar, 1 μm. (b) FRAP analysis of DeaD droplets *in vitro*. Error bars indicate s.d.; n = 6 condensates. Scale bar, 5 μm. (c) LLPS of purified recombinant DeaD in the presence of 1 μM mCherry in buffer containing 150 mM NaCl without PEG. Scale bar, 10 μm. (d) LLPS of purified recombinant DeaD with increasing concentrations of SerRS^wt^ or SerRS^T^ proteins in buffer containing 150 mM NaCl without PEG. Scale bar, 10 μm. (e) SerRS^T^ exhibited a comparable translocation pattern in DeaD-marked condensates formed with PEG. Data are mean ± s.d.; *P* values were calculated using two-tailed t-tests with Welch’s correction; n = 8 condensates, respectively. Scale bar, 10 μm. (e) SerRS^T^ exhibited a comparable translocation pattern in DeaD condensates formed with PEG. Data are mean ± s.d.; *P* values were calculated using two-tailed t-tests with Welch’s correction; n = 8 condensates. Scale bar, 10 μm.

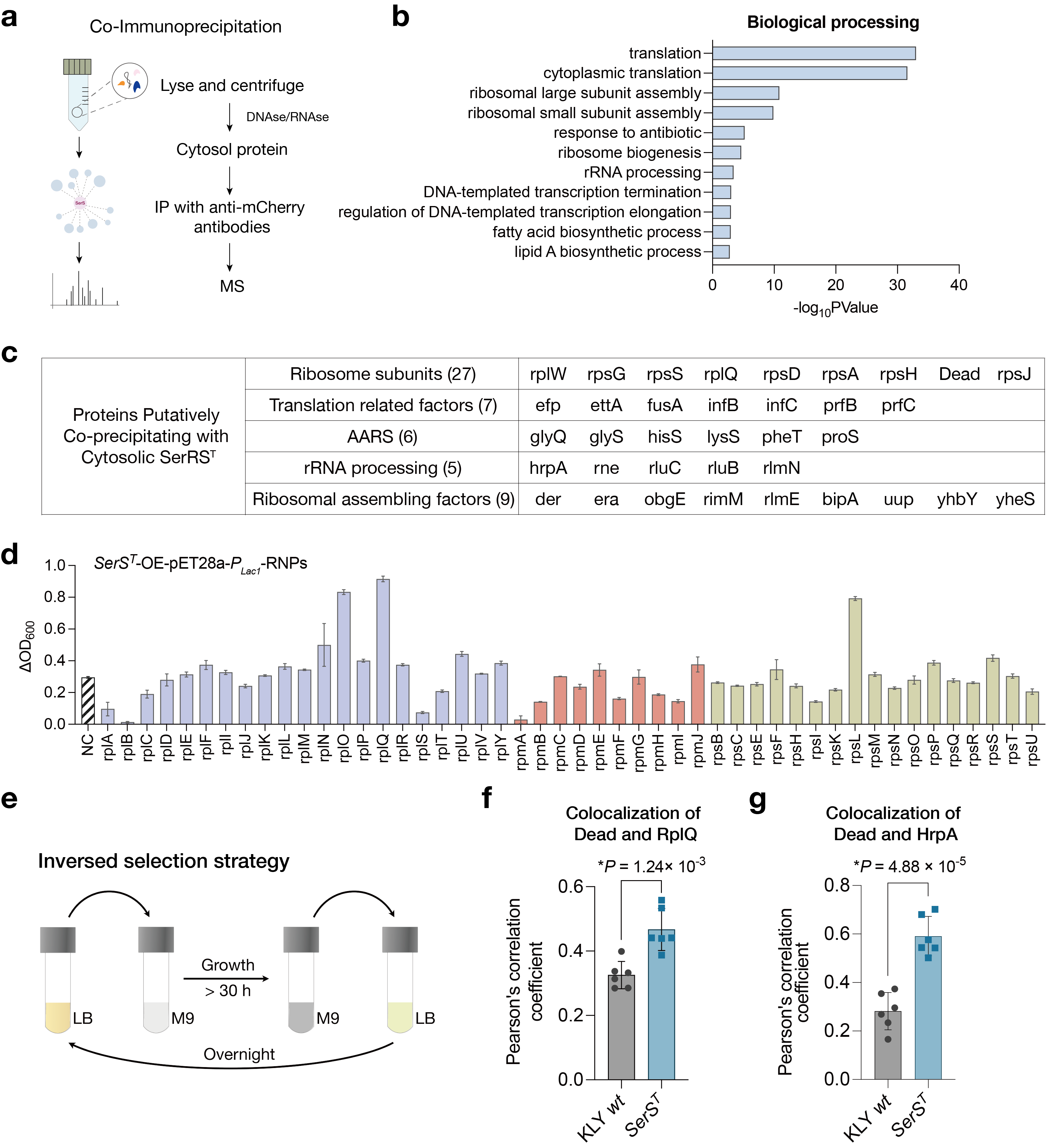

**Fig. S7 | Altered composition of condensates in the *SerS^T^* mutant compared to the KLY strain**

(a) Schematic of the experimental workflow, including cellular fractionation, immunoprecipitation, and LC-MS/MS analysis to identify SerRS-interacted proteins in the cytosol. (b) GO analysis of proteins immunoprecipitated from the SerRS^T^ versus SerRS^wt^ interactomes. The top 10 GO biological process annotations were shown as a bar chart, ranked by frequency (top to bottom), with the horizontal axis representing the -log10 *P* values of each term. (c) List of proteins identified by LC-MS/MS. (d) Effect of ribonucleoprotein (RNP) overexpression on the recovery of growth defects in the *SerS^T^* tolerant strain grown in LB medium. Purple and orange bars denote proteins of the 50S ribosomal subunit, while light green bars correspond to the 30S ribosomal subunit. (e) Schematic of the inversed selection experimental design. (f, g) Pearson’s correlation coefficients for colocalization of DeaD with RplQ (f), and DeaD with HrpA (g). Data are presented as the mean ± s.d.; *P* values were calculated using two-tailed t-tests.

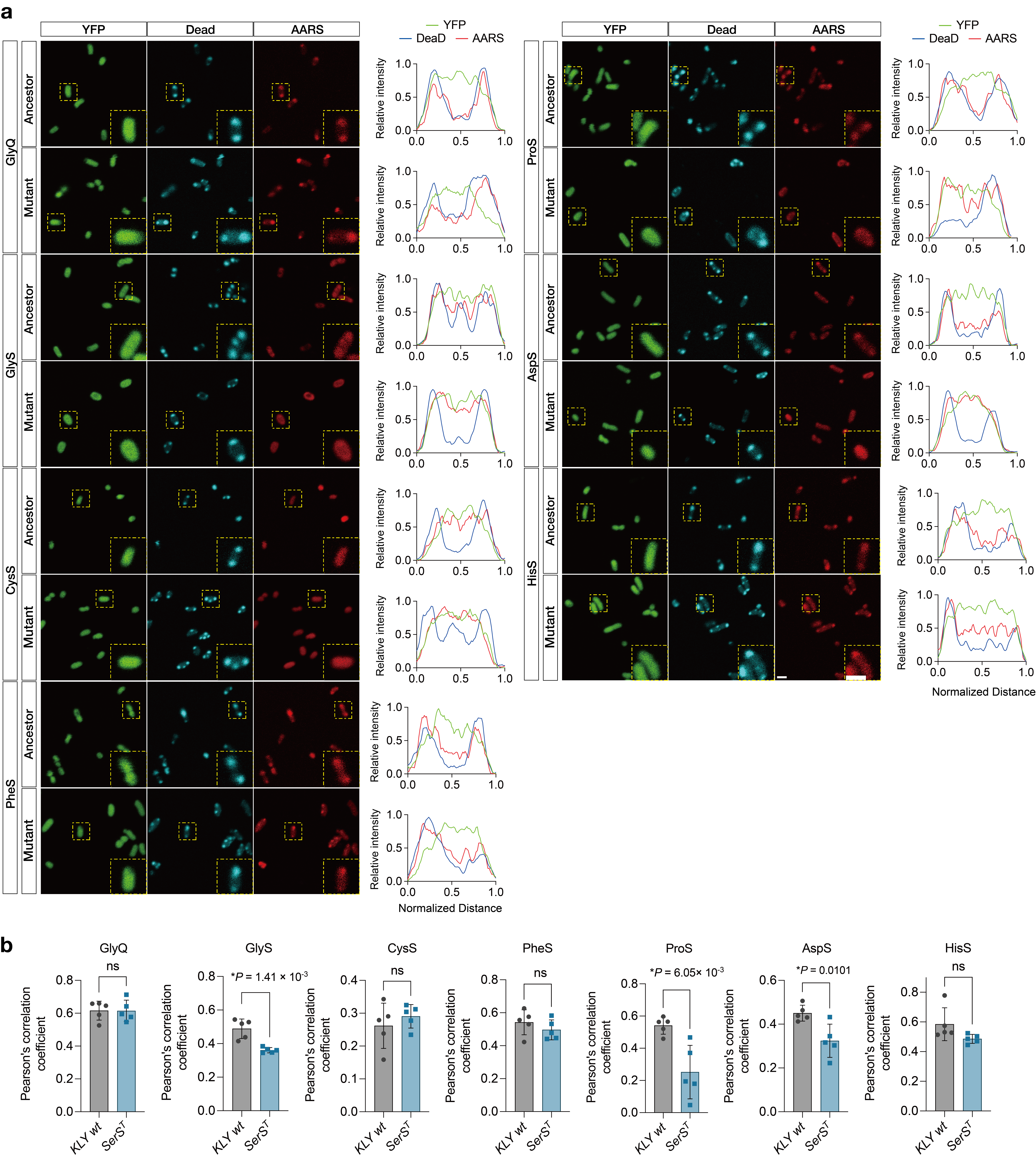

**Fig. S8 | Altered colocalization of aaRSs and DeaD-marked condensates.**

(a) Representative fluorescence images of mCherry-tagged aaRSs in cells expressing ECFP-tagged DeaD in the KLY *wt* and *SerS^T^* mutant cells. Inset: higher magnification of the yellow boxed area. Line scans show the related intensity profiles of aaRSs with DeaD. Scale bar, 2 μm. (b) Pearson’s correlation coefficients for colocalization analysis. Data are represented as mean ± s.d.; *P* values were calculated using two-tailed t-test.

**Table S1.** **Main mutations detected in whole-genome sequencing data and verified by Sanger sequencing**

| Strain | Locus | Amino acid substitution | Gene | Annotation |
| --- | --- | --- | --- | --- |
| KLY  R6E2a (*SerS^T^*) | 939,081 | G208D (GGT -> GAT) | *serS* | Seryl-tRNA synthetase |
| KLY  R15E2b | 939,066 | T203R (ACG -> AGG) | *serS* | Seryl-tRNA synthetase |
| KLY  R14E2c | 553,976 | P112L (CCG -> CTG) | *cysS* | cysteinyl-tRNA synthetase |
|  | 3,858,644 | M -> I (ATG -> ATA) | *cysE* | Serine acetyltransferase |
|  | 3,563,333 | N66K (AAT -> AAG) | *crp* | cAMP-activated transcriptional regulator |
| KLY  R11E2d | 939,303 | I282S (ATC -> AGC) | *serS* | Seryl-tRNA synthetase |

**Table S2. The plasmids used in this study**

| Name | Source |
| --- | --- |
| p15A-*P_tet_*-DeaD (Kan^R^) | This paper |
| p15A-*P_tet_*-DeaD-His (Kan^R^) | This paper |
| p15A-*P_tet_*-SerRS^wt^ (Kan^R^) | This paper |
| p15A-*P_tet_*-SerRS^wt^-mCherry (Kan^R^) | This paper |
| p15A-*P_tet_*-SerRS^T^ (Kan^R^) | This paper |
| p15A-*P_tet_*-SerRS^T^-mCherry (Kan^R^) | This paper |
| p15A-*P_tet_*-HisS-mCherry (Kan^R^) | This paper |
| p15A-*P_tet_*-GlyS-mCherry (Kan^R^) | This paper |
| p15A-*P_tet_*-GlyQ-mCherry (Kan^R^) | This paper |
| p15A-*P_tet_*-AspS-mCherry (Kan^R^) | This paper |
| p15A-*P_tet_*-ProS-mCherry (Kan^R^) | This paper |
| p15A-*P_tet_*-CysS-mCherry (Kan^R^) | This paper |
| p15A-*P_tet_*-PheS-mCherry (Kan^R^) | This paper |
| p15A-*P_tet_*-HrpA (Kan^R^) | This paper |
| p15A-*P_tet_*-HrpA-mCherry (Kan^R^) | This paper |
| p15A-*P_tet_*-SrmB-mCherry (Kan^R^) | This paper |
| p15A-*P_tet_*-RhlE-mCherry (Kan^R^) | This paper |
| p15A-*P_tet_*-ObgE-mCherry (Kan^R^) | This paper |
| pET28a-*P_lac1_*-DeaD (Amp^R^) | This paper |
| pET28a-*P_lac1_*-DeaD-ECFP (Amp^R^) | This paper |
| pET28a-*P_lac1_*-RBPs-mCherry (Kan^R^) ^a^ | This paper |
| p15A-PBAD-Cas9-PT5-Red𝛄𝛽𝛂 | Yi-xin Huo Provided (Beijing Institute of Technology) |
| PBAD-sgRNA-Donor ^b^ | Yi-xin Huo Provided (Beijing Institute of Technology) |

Note:

^a.^ all the ribosomal subunits were respectively overexpressed in this plasmid.

^b.^ PBAD-sgRNA-Donor containing different editing fragments.
